# A solid beta-sheet structure is formed at the surface of FUS liquid droplets during aging

**DOI:** 10.1101/2023.06.02.542764

**Authors:** Leonidas Emmanouilidis, Ettore Bartalucci, Yelena Kan, Mahdiye Ijavi, Maria Escura Pérez, Pavel Afanasyev, Daniel Boehringer, Johannes Zehnder, Sapun H. Parekh, Mischa Bonn, Thomas C. T. Michaels, Thomas Wiegand, Frédéric H.-T. Allain

## Abstract

Insights into liquid droplet formation via liquid-liquid phase separation and the subsequent liquid-to-solid phase transition are important for understanding cell dynamics, as well as a number of neurodegenerative disorders. We report here, using the example of the FUsed in Sarcoma (FUS) protein, an investigation of the liquid droplet maturation process combining solution- and solid-state NMR spectroscopy, Raman spectroscopy, and light and electron microscopies. Our study reveals that the surface of the droplets plays a critical role in this process. Indeed, when comparing a biphasic sample, in which liquid droplets are stabilized in an agarose matrix, with a pure monophasic condensed phase sample, we find that the liquid-droplet maturation kinetics is faster in the biphasic FUS sample, owing to the larger surface-to-volume ratio. In addition, using Raman spectroscopy, we observe structural differences upon liquid-droplet maturation between the inside and the surface of liquid droplets, which is of β-sheet content as revealed by solid-state NMR. This is detected very early on and increases over time. In agreement with these observations, a solid crust-like shell is visually seen by microaspiration experiments. After several months, electron microscopy reveals that the matured FUS droplets have converted into solid linear fibrils distinct from the fibril core of seeded fibrils reported previously, as arginine side-chains from the arginine-and-glycine-rich domain (RGG) motif are partially rigidified, highlighting the participation of this motif in the liquid-to-solid transition. In presence of RNA, this aging process is not taking place.

## Introduction

Research on the phase separation behavior of biomolecules has exploded in recent years as gradually more cellular functions are being discovered to rely on such phenomena [1], [2]. Many proteins will phase-separate from the aqueous environment to form an additional phase via a plethora of transient weak noncovalent interactions and may assemble into cellular membraneless organelles [3]–[5]. A now-commonly detected state is a liquid condensed phase, which enables rapid material exchange with the surrounding cyto- or nucleoplasm [6]. *In vitro*, this behavior can be reconstituted by the formation of liquid droplets. Interestingly, the liquid state of these entities, both *in vitro* and *in vivo,* may not be thermodynamically stable. Indeed, over time some of these liquid droplets transition into a less dynamic and often even solid-like state through a process known as maturation or aging [1], [7]. Since this solid state has been linked to various neurodegenerative diseases, it is of great interest to understand the mechanism of this transition and the associated loss of the dynamic nature of the liquid droplets. To date, no proposed model or mechanism posits how protein droplets transition to different states. In particular, which physical properties of the droplets allow for gradual solidification and how the atomic structure of the molecules influence the state of the matter are some of the key questions that remain elusive.

Here, we utilize the RNA-binding protein of FUS, which self assembles *in vivo* under stress to form stress granules in the cytoplasm [8]. FUS, like other RNA-binding proteins, has been previously shown to phase separate *in vitro*, and the resulting liquid droplets have been reported to rigidify over time, with disease-related mutations promoting significantly faster maturation than the wild type [9]–[11]. Furthermore, a short segment of the N-terminal unstructured half of the protein was shown to form amyloid fibrils after several days *in vitro* [12]–[14] though a complete molecular pathway of this liquid-to-solid transition (LST) is not known. Here, we investigate the liquid-to-solid transition of FUS droplets stabilized in agarose using a combination of (solution and solid-state) NMR spectroscopy, spatially resolved coherent Raman spectroscopy, electron microscopy and micropipette aspiration, and revealed the molecular pathway of the FUS droplet-maturation process.

## Results

### First days of FUS maturation studied by solution-state NMR

To study FUS-droplet maturation, we prepared several biphasic FUS N-terminal domain (NTD) samples (Fig 1a, residues 1-267) stabilized inside an agarose hydrogel as described previously [15]. FUS droplets were stable inside the hydrogel matrix for weeks. Proteins inside FUS liquid droplets diffuse at least 100 times slower compared to outside [16]. This difference enables us to observe and quantify the fraction of protein within liquid droplets (henceforth condensed phase) using Diffusion Ordered Spectroscopy (DOSY) NMR experiments at maximum gradient strength [15]. Conversely, the spectra at low gradient strength report on both the condensed (liquid droplets) and dilute (dissolved proteins) phases. Therefore, the difference between the two, enables us to quantify the amount of protein in the dilute phase.

**Figure 1:**
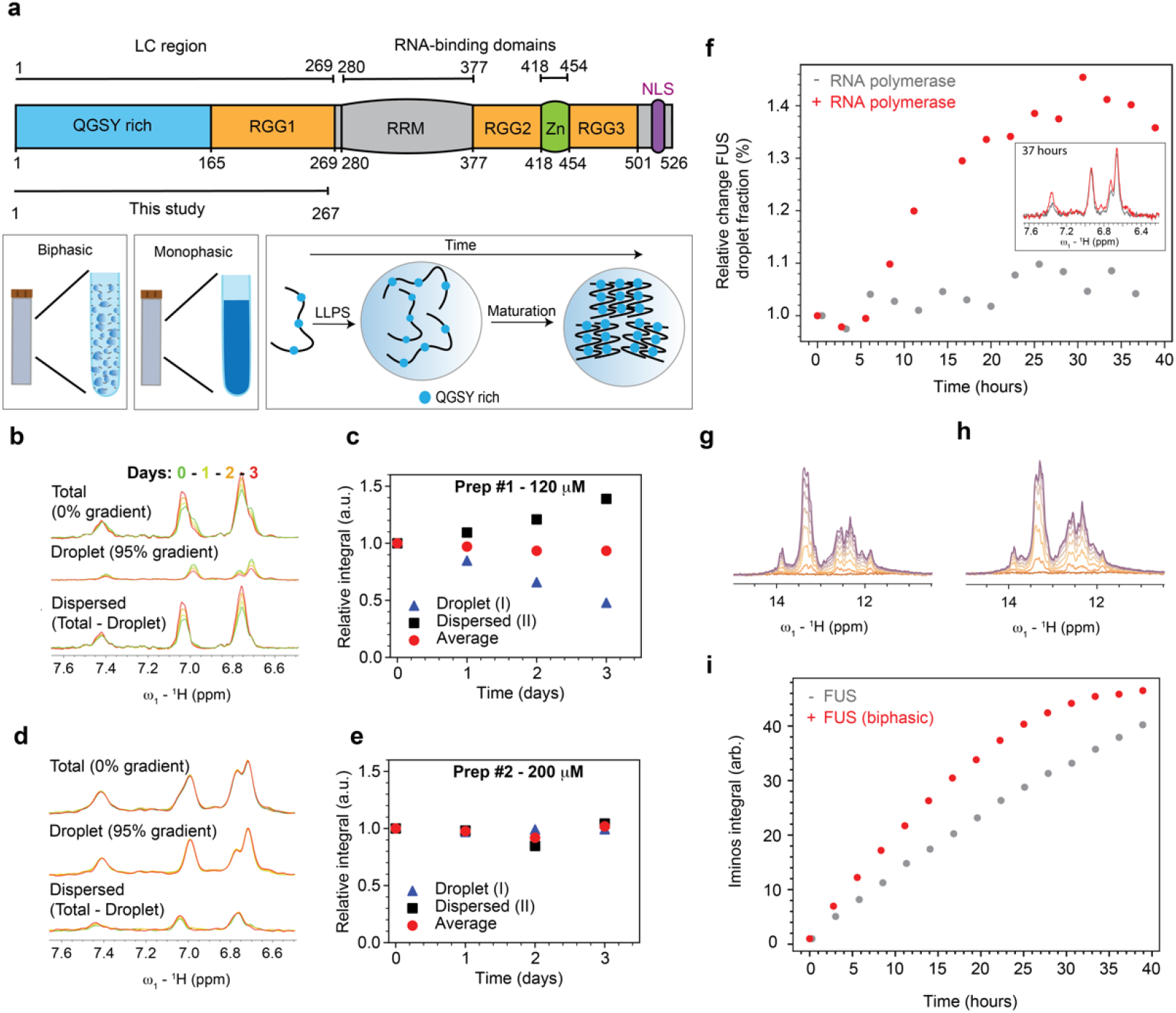
Evolution of FUS phase fraction over time studied by solution NMR. a) Schematic representation of FUS protein domain organization, LC = low complexity, RRM = RNA recognition motif, Zn = zinc finger, NLS = nuclear localization signal. Definition of biphasic and monophasic are also shown, biphasic contains liquid droplets while monophasic a single continuous dense phase as result of sentimentation of droplets. Maturation process is the transition from liquid droplets to less dynamic immobile species. b,c) Overlaid DOSY spectra at different time points at 120 μM (b) and the corresponding integrals for direct comparison (c). d,e). Overlaid DOSY spectra at different time points at 200 μM (d) and the corresponding integrals for direct comparison (c). At zero gradient strength the total population of FUS is visible, while at maximum gradient only the droplet phase. The difference between the two reports only the dispersed fraction. The two panels represent the two distinct modes of behaviour observed in this study. f) Relative FUS droplet fraction change within 40 hours in presence and absence of RNA transcription, with the corresponding ^1^H NMR spectra shown in the inlet. g,h) Overlay of 1H NMR spectra at different time points focused on the imino region in absence (g) and presence (h) of FUS liquid droplets, orange = 0 h progressively to magenta = 40 h. i) RNA iminos integral progression over time indicating the faster kinetics of the biphasic sample.

We measured a series of DOSY experiments over three days from different FUS preparations and at various protein concentrations (120 – 400 μM). Surprisingly, we observed two types of maturation behavior. In type 1 (two cases, corresponding to the lowest concentrations tested #1 – 120 μM and #3 – 180 μM), we measured a decrease of the NMR signal corresponding to the condensed phase and a simultaneous increase of the dilute phase fraction (Fig 1b, Fig S1a). The average of the two normalized intensities remained close to one over time, suggesting that part of the droplet phase gradually diffuses faster and possibly dissolves into the dispersed phase (Fig 1c). However, since we do not observe any obvious turbidity change within the NMR tube, we presume that the droplets did not dissolve. In type 2, at higher initial FUS concentrations (#2 - 200 μM and #4 – 400 μM), we observed no significant evolution of the fractions over the first three days (Fig 1d,e, Fig S1b). To rationalize these observations, we developed a simple theoretical model that explores the relationship between FUS aggregation kinetics and phase separation using recently established theories for chemical reactions in coexisting phases at phase equilibrium [17], [18] (see Fig. S2a and Materials and Methods for details). Using this model, we explored how aggregation kinetics couple to the physical properties of droplets, in particular changes in solvent volume fraction, under two different regimes, either when (i) aggregate-solvent interactions are favoured over monomer-solvent interactions and (ii) vice-versa. Interestingly, in the relevant first scenario, the model predicts a protein concentration-dependent solvent flux into the droplet phase (Fig. S2b,c) which would explain the increase of diffusivity as aggregates accumulate inside the droplets as shown by NMR (see Supplementary materials section). Our results are consistent with a previous study which shows how aggregation couples to changes in droplet volume over time [19]. While these simulations show clear changes of solvent flux into the droplets that are consistent with our NMR data, certainly other factors may also contribute to the observed NMR signal changes (Fig S3).

### FUS maturation is suppressed in the presence of RNA

As FUS is known to play a role in RNA transcription [20]–[22], we studied if transcription could influence FUS maturation *in vitro*. To this end, we prepared *in vitro* transcription reactions as previously described [23] and quantified the reaction velocities in the absence and presence of agarose-stabilized FUS droplets. Since no droplets are formed at 37°C, the experiments were performed at 25°C. Lowering the temperature resulted in slower reactions that reached saturation only after two days. This provided a sufficiently long time window to follow the velocity of transcription, but also the FUS droplet maturation process. Interestingly, we observed a strong, irreversible increase (up to 40%) of the condensed form of FUS (Fig 1f) during the transcription reaction over 40 hours. Without transcription, the droplet fraction of FUS is stable over time (Fig 1f). As reported previously, low concentrations of RNA enhance droplet formation of RNA-binding proteins, while more RNA dissolves them [24]. Hence, observation of a large increase of the fraction of FUS in the condensed phase during the reaction indicates that the transcribed RNA interacts with FUS and further promotes its phase separation. Since the level of FUS in the droplet form is maintained for several days, this suggests that RNA promotes FUS liquid condensation over droplet maturation. Another interesting observation could be made when comparing the solution-state NMR RNA imino signals during the time course of the reaction in the presence and absence of FUS. We could measure in the biphasic sample of FUS a strong increase in the amount of produced RNA due to an increase in the initial transcription speed (Fig 1g,h,i). While further experiments are required to elucidate the cause of this effect, the increased velocity could be attributed to crowding effects or interactions between FUS and reaction components that favor the polymerase kinetics.

### FUS maturation monitored by real-time solid-state NMR

To directly observe the maturation of liquid droplets and the formation of potential solid fibril species, we turned to solid-state NMR, which can detect immobilized species and was previously applied successfully to determine the structure of the FUS fibril core (residues 39 to 95) [12] and to study the maturation kinetics of a monophasic sample of FUS (1-163) in the condensed phase [14] (Table S1).

We tracked droplet maturation of our longer ^13^C,^15^N-labelled FUS-NTD (1-267) sample in an agarose matrix (∼8 mg of protein in the NMR rotor) over two days by real-time refocused Insensitive Nuclei Enhancement by Polarization Transfer (INEPT) and cross-polarization (CP) NMR spectra, which allow the detection of soluble (highly mobile) and immobilized species, respectively [25]–[27]. Unlike the DOSY experiment, INEPT could not distinguish between NMR signals originating from the dilute and condensed phases. Nonetheless, we observed only a weak signal-intensity decrease (∼10%) in the INEPT spectra over the first two days of maturation, followed by magic-angle spinning (MAS) (Fig 2a, Fig S4a). This agrees with our solution-state NMR observations, where the net protein signals (the sum of the dilute and liquid-droplet phase) remained nearly constant for the initial three days of maturation (Fig 1c,e). By contrast, over a longer period (37 and 73 days), we detected an increase of the INEPT signal combined with a significant narrowing of the resonances due to an increase in the molecular-tumbling rate (Fig S4b). Turning to CP, we detected a signal already 4 h after rotor filling. The intensity of the CP signal increased over the course of two days, followed by a further increase within the next ∼35 days, and no further changes after 73 days (Fig 2a, Fig S5a,b). Note that the comparison of 1D CP spectra recorded in different measurement slots has an associated uncertainty in the peak intensity of roughly ±10%. We see no obvious chemical-shift perturbations during the first two days of maturation, indicating the gradual growth of a solid material already present four hours after droplet formation (Fig S6). Quantitatively, half of the final CP signal observed after two days was already present after 4 hours in the first CP spectrum (Fig 2a), indicating that it is produced relatively quickly after droplet formation and spinning for the MAS experiment. The initial formation of solid material thus occurs very rapidly, and only the subsequent increase is linear in time, but is not associated with a corresponding decrease in the INEPT signal as observed, for instance, in real-time solid-state NMR studies on protein aggregates [27]. This discrepancy between INEPT- and CP-observed protein signal during FUS maturation has been previously reported and points to the potential presence of an invisible population being converted into solid over time [14]. Comparing the ^13^C CP spectra after two days of real-time MAS NMR and after 37 and 73 days revealed some spectral differences, indicating the formation of structurally distinct species after a longer maturation period (Fig S5a,b).

**Figure 2:**
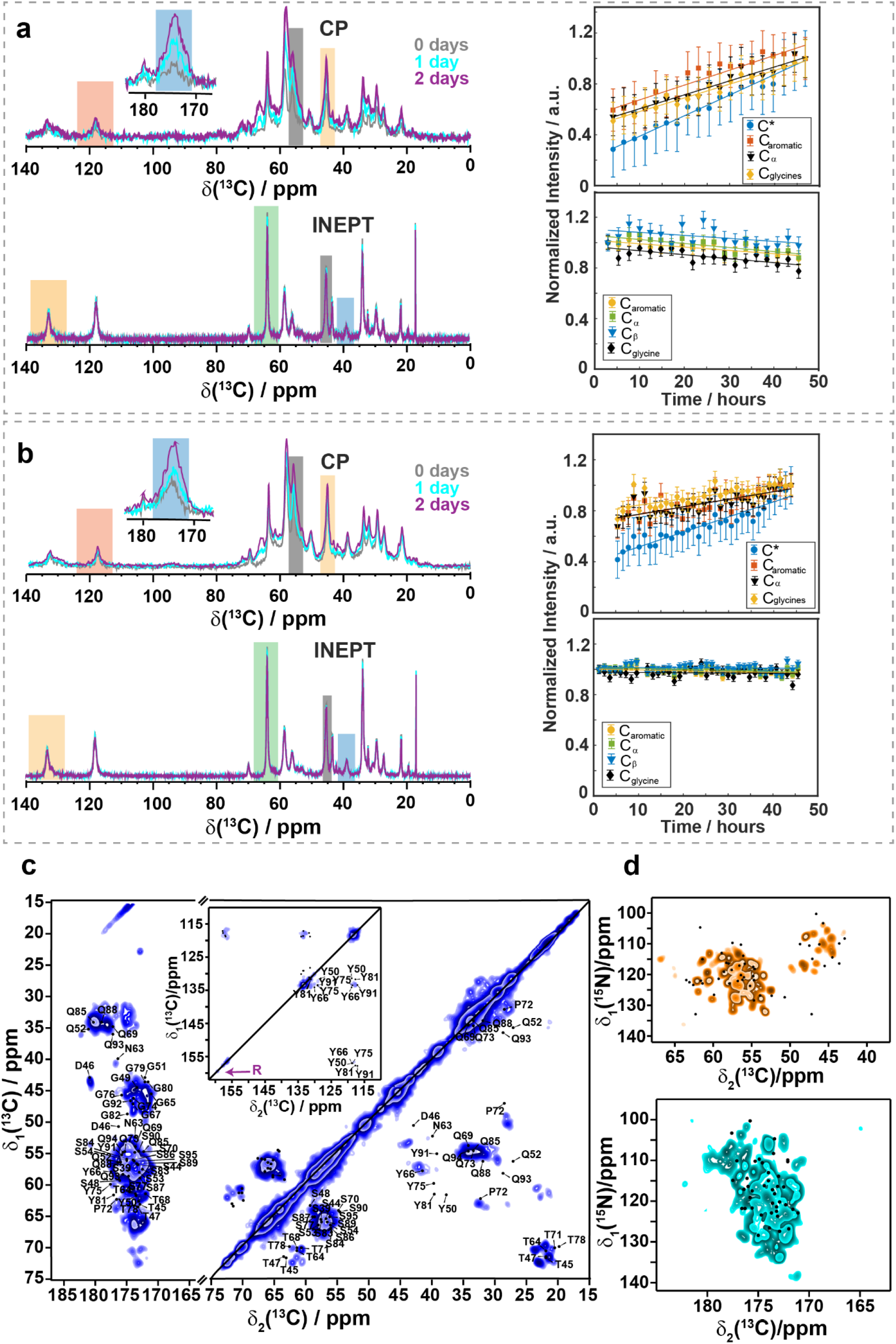
a) Time-dependent ^13^C,^1^H CP-MAS and ^13^C,^1^H INEPT spectra of biphasic FUS (left), as well as time-dependent intensity changes during two days of maturation for several selected resonances (right, the spectral regions used are highlighted by colored rectangles in the spectra). The intensity of the CP spectrum recorded after two days was normalized to one. A linear regression is shown (straight lines) with slopes (units of 1/days) of 0.389 (C*), 0.281 (C^aromatic^), 0.264 (C^α^) and 0.259 (C^α,glycines^) for the CP spectra and -0.062 (C^aromatic^), -0.077 (C^β^), -0.058 (C^α^) and -0.074 (C^α,glycines^) for the INEPT spectra. b) Time-dependent ^13^C,^1^H CP-MAS and ^13^C,^1^H INEPT spectra of monophasic FUS (left), as well as time-dependent intensity changes during two days of maturation for several selected resonances (right, the spectral regions used are highlighted by coloured rectangles in the spectra). The intensity of the CP spectrum recorded after 2 days was normalized to one. A linear regression is shown (straight lines) with slopes (units of 1/days) of 0.283 (C*), 0.144 (C^aromatic^), 0.142 (C^α^) and 0.120 (C^α,glycines^) for the CP spectra and -0.014 (C^aromatic^), -0.017 (C^β^), -0.005 (Cα) and -0.005 (C^α,glycines^) for the INEPT spectra. c) ^13^C-^13^C 20 ms DARR spectrum (left) of maturated monophasic FUS (68 days). The inset shows the aromatic region. A weak arginine-side-chain resonance is detected on the diagonal, highlighted by a purple arrow. The assigned peaks are back-predicted from [12]. d) ^15^N-^13^C NCA and NCO spectra of matured monophasic FUS (114 days). The dots represent peaks back-predicted from [12]. All spectra were recorded at 20.0 T and 17 kHz MAS.

Moreover, a visual inspection of the agarose-stabilized droplets after six months revealed the presence of suspended white particles. After extraction, we characterized the nature of these particles by electron microscopy and observed fibrils and fibril bundles (Fig S5c). While further studies will be required to identify the atomic details of these FUS fibrils, our NMR data already indicate a different fibril fold compared to the shorter construct obtained upon seeding (1-163, see below) [12].

### Different maturation rates in biphasic and monophasic samples

Another key difference from previous studies on FUS fibrils [14] is our sample preparation protocol. In our study, FUS droplets were stabilized inside an agarose hydrogel matrix, where they could mature as suspended droplets. In previous studies, droplets were centrifuged, resulting in a single bulk condensed phase from which the fibrils were formed and harvested. Liquid droplets possess a much larger combined surface compared to a bulk phase. To gain insight into the role of surfaces in maturation, we also tracked the development of solid-state NMR signals over time in a sample containing a single bulk condensed phase to compare it to the sample of liquid droplets. We used around 10 mg of protein sample in the 3.2 mm rotor for the bulk condensed phase (in the following denoted as monophasic sample).

Similar to FUS droplets in the biphasic sample, the monophasic sample did not yield any significant change in the INEPT signal intensities over two days (Fig 2b). On longer time scales, the relative intensity increase in the INEPT spectra was small compared to the biphasic sample (Fig S7a,b). Similarly to the biphasic sample, some immobilized species were already detected in the first CP measurement taken after 4 h (Fig 2b). However, two decisive differences were observed in the maturation process between the two samples. The first one is that at the beginning of the maturation period, the CP spectra are very similar (Fig S8a), but after two months of maturation, they differ concerning peak positions and peak intensities (Fig S8b and the spectral regions highlighted therein). This is in line with the second major difference, namely changes in the maturation rates between the two samples. A comparison of the slope from the integrated resonances over time revealed that the biphasic sample matured approximately two times faster than the monophasic one (Fig 2a,b, Fig S10a,b). For the monophasic sample, this is surprising because it contains more protein mass (∼10 mg of FUS protein in a volume of 46 μL), while the biphasic sample contains less protein (∼8 mg). While the biphasic sample shows a clear linear increase of the integrated CP resonances over time (Fig S10a), some deviations from this behavior are observed for the monophasic sample (Fig S10b).

Overall, this higher maturation rate found in the biphasic sample points to a clear role of the liquid-droplet surface. As mentioned previously [15], the surfacearea-to-volume ratio is substantially larger in the biphasic sample, where droplets are stabilized, than in the monophasic bulk condensed phase. Since the two samples studied here also differ in agarose content, we investigated if the droplet surface would be the primary site of the maturation process.

### FUS maturation results in structural differences between the surface of the droplets and the interior as probed by Raman spectroscopy

We exploited Raman spectroscopy to study the structural differences at the surface of droplets. This method has the advantage of combining spatial resolution with protein secondary structure detection. By diluting FUS inside the agarose hydrogel, micrometer-sized droplets were formed, and we measured Raman spectra in distinct areas of month-old droplets (Fig 3a,b). Analysis of the recorded spectra in the amide I region, which is sensitive to the protein secondary structure, revealed clear differences between the inside and the surface of the droplet. Further analysis of the spectra by segmenting the droplet into concentric rings showed that droplet possessed a core-shell like structure: The interior of the droplets was largely homogeneous, as the spectrum only changed very close to the droplet surface (Fig 3c). Interestingly, normalized spectra revealed a more intense tyrosine peak at 1618 cm^-1^ at the surface (Fig 3c). As this tyrosine peak is highly sensitive to hydrogen bonding, a more intense peak corresponds to tyrosines participating in a stronger hydrogen bond network, as expected from fibrils containing tyrosines in their cores [28]. The importance of tyrosine residues in liquid-droplet maturation is further underlined by the rigidification of their side-chains in the final fibrils suggesting π-π stacking as shown by their appearance in the solid-state NMR CP spectra (see below).

**Figure 3:**
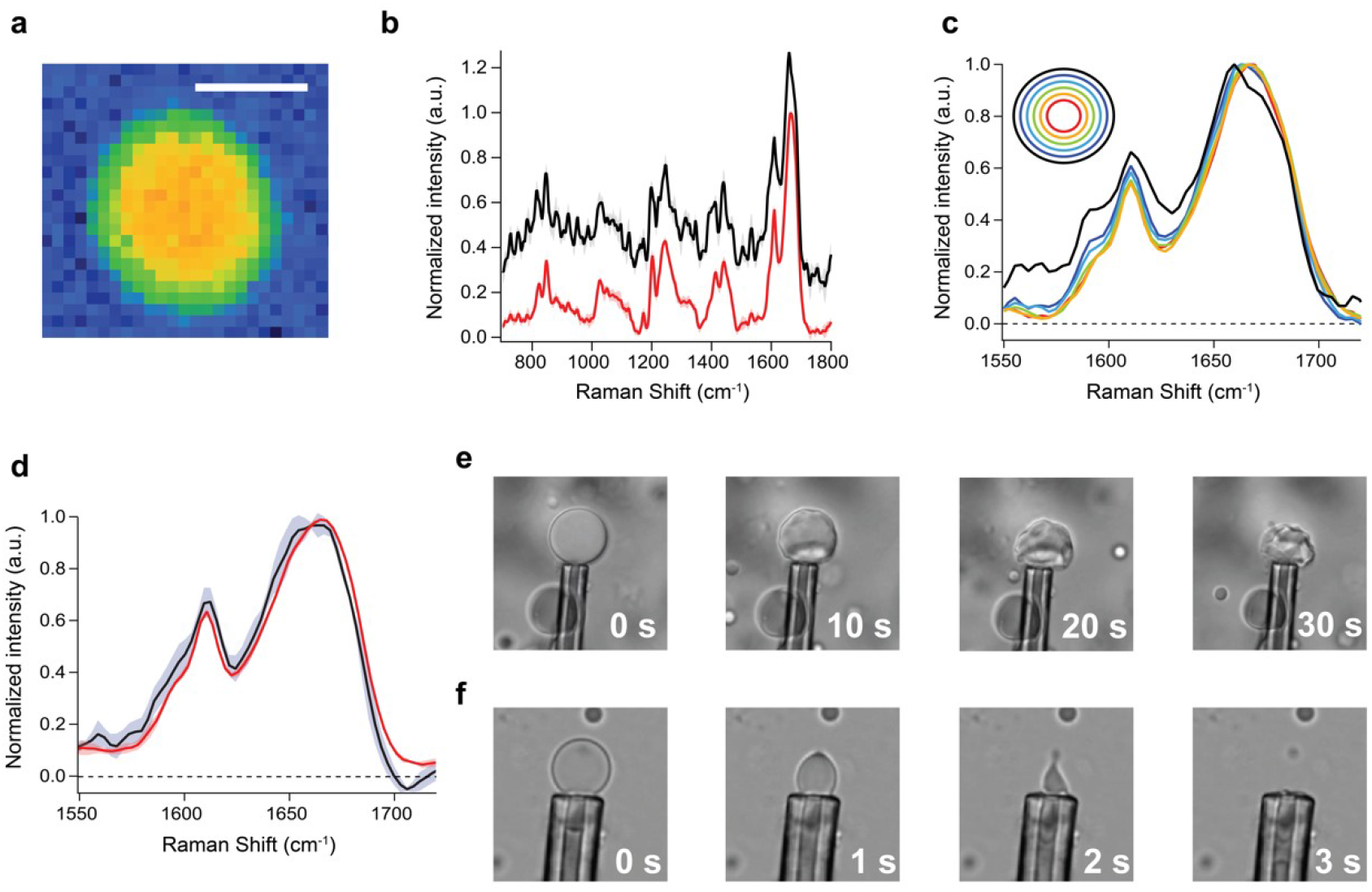
a) Image of 2D scanned FUS-NTD (residues 1-267) with the pixel contrast given by integrated intensity of amide I band. Scale bar : 4 μm. b) Fingerprint region normalized averaged Raman-like spectra of one month old droplets’ internal region (red) and border region (black), shaded areas show the deviations between different samples (N=5). c) Comparison amide I band spectra of the concentrical rings regions of the droplets. In black is shown the spectrum corresponding to the droplet surface. d) Comparison amide I band spectra of the internal region (red) and border region (black) of fresh droplets, shaded areas show the deviations between different samples (N=5). (e,f) Microaspiration of matured FUS droplets in absence (d) and presence (e) of polyU RNA. The droplet can be completely aspirated only in presence of RNA, while in absence a solid shell-like structure is revealed on the periphery of the droplet.

The spectrum at the surface of droplets exhibited a significantly narrowed peak at 1665 cm^-1^ which is absent in the inside of the droplet (Fig 3c). Taken together, the spectra at the surface and inside the droplets clearly reveals spectral differences pointing to significant structural changes occurring on the droplet surface upon maturation, while no changes are detected in the fresh droplets (Fig 3d).

Next, we tested the material properties by physically probing the droplets. Specifically, we aspirated material from a month-old matured droplet to determine the material state and continuity. While we could aspirate the liquid interior of the droplet, we observed that the periphery is made of a solid crust-like material that could not be aspirated and therefore collapsed (Fig 3e, Suppl. Video 1). Interestingly, the subsequent addition of polyU RNA resulted in shell-less droplets, which can be aspirated ten times faster at the same pressure and with similar sized droplets (Fig 3f, Suppl. Video 2). Thus, it appears that the RNA can liquefy the already-rigid droplet shell, most likely by interacting non-specifically with the RGG segment of the protein. This is in alignment with previous reports showing that RNA can preserve the liquid state of the FUS droplets and decrease the viscosity of the RGG-containing LAF-1 droplets [29], [30]. This also agrees with our in vitro transcription experiment, where RNA stabilizes FUS droplets and prevents their maturation (Fig 1).

Since the bulk of the aged droplets can be aspirated, we conclude that it remains as a liquid, confirming our solution-state NMR data and our theoretical model (Fig 1 and Fig S2). Conversely, the surface appears solid-like, suggesting that it is a more structured material. The liquid-crust contrast between the bulk and interface of the droplet in aspiration agrees with the differences we see in the Raman spectra, which clearly points to a different FUS structure inside and at the surface of the droplet. In combination with solid-state NMR data, where we observe structural changes during maturation, leading to an increase of β-sheet content (*vide infra*), we can now propose that the crust-like material observed for matured droplet is likely to be primarily of β-sheet nature.

### FUS fibril secondary structure studied by solid-state NMR

To characterize the molecular structure of the solid material formed during the liquid-to-solid transition, we recorded 2D ^13^C-^13^C Dipolar Assisted Rotational Resonance (DARR) spectra [31], [32] and ^15^N,^13^C NCA and NCO spectra of the matured samples of the bi- and monophasic samples [33]. Fig 2c shows the 2D DARR spectrum for the 68 days matured monophasic FUS sample. The spectrum is well resolved with ^13^C line widths of ∼1 ppm (∼240 Hz), pointing to rather homogeneous fibrils formed in the phase transition process. ^13^C Cα and Cβ chemical-shift values are sensitive reporters for the secondary structure [34], which has been employed in structure calculations of amyloid fibrils [35]. For the spectrally well-resolved threonine and serine residues located in β-sheets, α-helices, and loops, we have plotted the statistic distribution of ^13^C Cα/Cβ chemical-shift values from probability density functions [36]–[38] (for more details, see Supplementary materials section, and Fig S11). This analysis confirms the formation of β-sheets in matured samples, in agreement with previous studies of shorter FUS LC constructs (residues 1-163) [14]. Furthermore, we inspected the 1D ^13^C CP spectrum of the monophasic FUS-NTD obtained after two days of maturation (Fig. S11a,b) and observed the presence of threonine Cα/Cβ resonances characteristic for β-sheet secondary structures (∼61.2/ 72.3 ppm for the Cα and Cβ resonances) showing that β-sheet already appeared within the first two days of maturation. We thus plotted on the 1D ^13^C CP spectrum the threonine ^13^Cβ averaged chemical-shift values for the various secondary-structure types (Fig S11b) [36]. Indeed, the high-frequency shifted threonine Cβ resonance falls within the β-sheet region. Further comparison of the 1D spectrum with the well-resolved threonine shifts in the 2D DARR of the sample after 68 days of maturation shows a clear match of the resonances (Fig S11c). Together with the Raman experiments, the fact that the biphasic and monophasic samples have almost identical 1D CP (Fig. S8) allow us to propose that the initial β-sheet formation takes place at the surface of liquid droplets and helps in the formation of the crust observed during microaspiration.

Comparing our DARR spectrum to the one published previously on shorter FUS constructs reveals several differences (Fig S12) [14]. First, a variety of additional resonances is observed in our case (for instance, in the threonine region, orange boxes in Fig S13), together with noticeable chemical-shift perturbations for some well-isolated resonances (purple boxes in Fig S12). Second, we could also detect signals from arginine side-chains in a ^15^N CP MAS spectrum (Fig S13a), which apparently also rigidify to some extent after the phase transition process. This is notable as arginine residues are only present in the RGG domain (absent in FUS LC), which suggests its engagement in the phase-separation process. The importance of the RGG domains in liquid-liquid phase separation (LLPS) has been reported recently in the context of the full-length protein [39], and an indication for the role of the RGG domain in LLPS of FUS-NTD is obtained by the solid-state NMR spectra presented herein. The matured FUS fibrils also show an intense CP signal for rigidified aromatic tyrosine side-chains (Fig S13b,c), which are efficiently immobilized, most likely due to π-π stacking interactions or hydrogen bonds as indicated also in the Raman spectra. Interestingly, and as reported also previously [14], the solid material formed via phase separation presents clear structural differences to the FUS fibrils grown from fibril seeds (residues 1-214), for which the fibril core (residues 39-95) was structurally solved by solid-state NMR [12]. This is evident by the differences in chemical shifts between the LLPS-induced fibril state and the fibril core reported previously (see the back-predicted shifts for the FUS fibril core only plotted on the 2D DARR spectrum in Fig 2c and the 2D NCA and NCO spectra in Fig S14). Similar conclusions can be drawn from the highly resolved heteronuclear NCA and NCO spectra shown in Fig 2d, which possibly also open the way for a resonance assignment in the future. Unfortunately, due to the low signal-to-noise ratio (see Fig. S15 for the 2D DARR spectrum), such an analysis was not be possible with the current biphasic sample.

## Discussion

Even though we can describe the maturation of liquid droplets both at the macroscopic (solidification) and atomic (fibrilization) levels, we lack information on how these two aspects are connected and therefore on the mechanism of the liquid-to-solid transition (Fig. 4a). Here, we propose a mechanism of the liquid to solid transition that incorporates our experimental observations (Fig. 4b). Our data allowed us to better understand a previously reported NMR-invisible population of FUS [14] that we detect as well with our solid-state INEPT decrease (Fig 2a, Fig S4a). This spectroscopically invisible FUS may be positioned at the surface of the droplets and progressively be converted to a solid crust where it becomes visible again (Fig. 4b). This state escapes detection by solid-state NMR due to its motion rate that may cause an inefficient heteronuclear polarization transfer by INEPT, as well as by CP experiments [40]. The role of the surface in maturation is supported by the faster rate of fibrilization of the biphasic sample compared to the monophasic revealed by solid-state NMR and different structural composition detected on the surface of the droplets compared to the droplet interior by Raman spectroscopy. The partial rigidification of arginine side-chains in the RGG1 domain might indicate a different fibril core segment than reported previously [12], [41], which is also supported by significant chemical-shift differences between fibrils obtained from liquid-droplet maturation and seeded fibrils. Alternatively, the partially rigidified arginine could correspond to surface-immobilized side-chains, as proposed by the coarse-grained model of Garaizar and coworkers [42]. Subsequently, the concentration of the fibrils increases on the surface, likely due to a difference in surface tension [43]. During this process, water enters the droplets to balance the equilibrium between the two phases. The additional water decreases the viscosity inside the droplets; hence the DOSY NMR signal for the fast-diffusing FUS increases. Ultimately, this process results in a solid shell on the periphery that macroscopically indicates the passage from a liquid to a partially solid phase. This barrier between the dilute phase and the inside bulk droplet phase prevents the exchange of protein material, resulting in the observed limited recovery after photobleaching [44].

**Figure 4:**
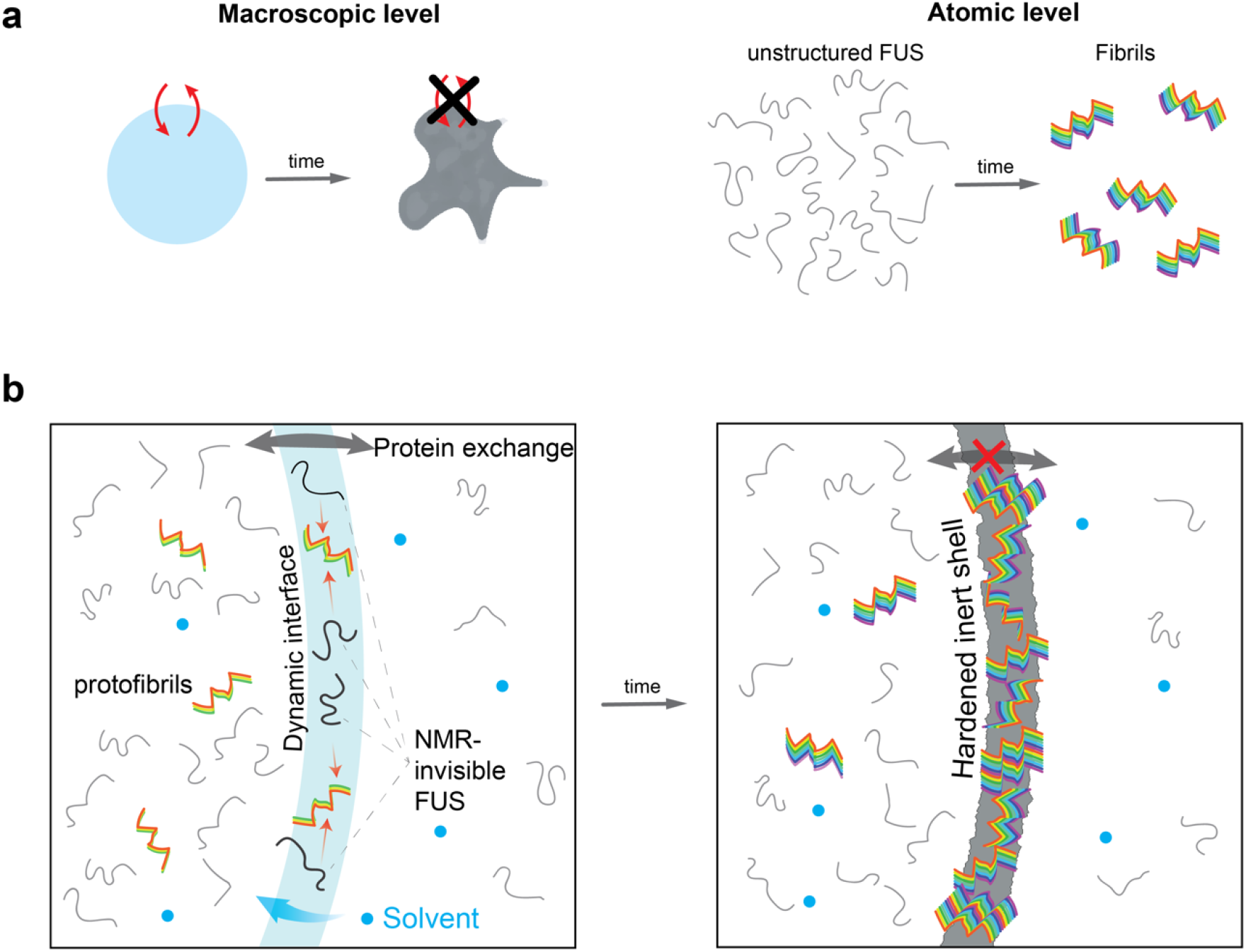
Current and proposed models for FUS maturation mechanism. a) Current view of maturation on macroscopic and atomic level. b) Proposed model. Fresh droplets have highly dynamic interface with the solvent allowing protein exchange between phases. While droplets are initially formed, a fraction of the protein instantaneously adopts a protofibrilar fold which is present both in the bulk and on the surface of the droplets. We here propose the existence of a FUS population on the periphery, which due to the unique environment of the interface escapes detection from conventional INEPT NMR experiments. Over time, this population is gradually incorporated into the fibrils of the surface. During the fibrilization process the hydrophobic nature of the droplet decreases as the hydrophobic groups of the protein are protected in the fibril core. This increases the flux of solvent molecules inside the droplet causing the diffusivity of the remaining monomers to increase. Eventually, once equilibrium is established all surface is converted to fibrillar species resulting in the inert hardened shelled observed which hinders material exchange between the phases.

The critical role of the droplet surface in the liquid-to-solid transition is only starting to emerge, as it was modeled recently [42], [45]. Our work presents spectroscopic evidence that maturation occurs at the surface of liquid droplets. This is in agreement with previous reports of Thioflavin T accumulation and high Förster resonance energy transfer (FRET) efficiency at the coacervates surface of TDP-43 and α-synuclein, respectively [46], [47]. Thus it creates an additional cellular sub-compartment to associate molecular functions or exploit as therapeutic target. Even though the concept of reactive liquid-liquid interfaces is not new in chemistry, where interfacial polymerization is utilized to obtain crystalline needles [48], [49], very few examples in biology have been reported. The synthesis of pre-ribosomal RNA occurs at the interface between two of the three phases that comprise the nucleolus [50]– [52]. More recently, anisosomes were discovered in cells described as liquid spherical shells made of the RNA-binding protein TDP-43. Although both the droplet interior and the surrounding shell exhibited liquid properties, the latter was much more dense and functioned as a selective barrier [53]. This architecture of a dense shell resembles our macroscopic observation of a solid-like droplet periphery, and future studies should reveal if it also shares the same function. Intriguingly, we report the sub-compartment of the droplet shell can be dissolved by RNA, which may indicate a specific function of the shell to orient charged side-chains, i.e. arginines, as predicted [42]. Collectively, the inhomogeneity of the liquid droplets and the actual functional role of their surface indicate that the droplets are the means to form this important sub-compartment, rather than the actual bulk condensed phase.

Maturation of FUS has been linked with disease evolution, although a debate exists. FUS disease mutation G156E matures significantly faster than the wild type [9]. If indeed the matured species causes the disease, then for a therapeutic approach, it is mandatory not to perturb the stress granules but rather to prevent disease-relevant maturation. Therefore, instead of designing very specific molecules, we could repurpose already known molecules that partition into hydrophobic interfaces acting as wetting agents and ultimately interfere with maturation.

## Materials and Methods

### Protein Expression and Purification.

FUS NTD (1-267) was expressed and purified under denaturing conditions as reported previously[15]. Double uniformly labeled protein (^15^N and ^13^C) for solid-state NMR experiments was expressed in M9 medium with ^15^N ammonium chloride and ^13^C glucose as the nitrogen and carbon source, respectively. The final stock concentration was 10 mM, unless otherwise stated, in 6 M urea buffer (50 mM Hepes, 150 mM NaCl, 6 M urea, pH 7.5).

### Solution-state NMR

Solution-state NMR samples were prepared as described previously[15]. Briefly, protein stock was diluted to the desired concentration using hot agarose buffer (30 mM Hepes, 200 mM, 0.5% wt/v agarose (ThermoFisher), pH 7.3). The sample was transferred to 3 mm NMR tube where gradually temperature dropped to room temperature (25 °C), causing agarose hydrogel and FUS liquid droplet formation.

All solution-state NMR experiments were recorded at 25 °C using the following Bruker spectrometers with *z*-axis pulsed field gradients: Avance III at 750 MHz proton frequency with a PATXI room-temperature probe (prep #1, #5), Avance NEO at 500 MHz equipped with a CPQCI cryogenic probe (prep #2), Avance III HD at 600 MHz (prep #3) and Avance NEO at 700-MHz proton frequency equipped with a CPTCI cryogenic probes. All spectra were processed with Topspin 3.2 (Bruker Biospin).

A standard pulse sequence (stebpgp1s19 from Topspin 3.2, Bruker Biospin) was used for diffusion experiments. In total, 4,096 points with 32 scans were recorded in the proton dimension for each dimension with variable diffusion gradient strength ranging between 2 and 95% in various steps. The following parameters were used: diffusion time (*Δ*) 0.05 or 0.1 s, gradient pulse (*δ*) 10 ms, smoothed rectangular-shaped gradients SMSQ10.100, relaxation delay (d1) 4s.

### *In vitro* transcription reactions.

The in vitro transcription reactions were designed according to the previously published protocol [23] with additional 0.5% wt/v agarose, if stated. The RNA transcribed is an intronic splicing regulator (downstream control sequence – DCS) from c-src gene [54]. Template DNA is cloned in a PTX1 [55] vector which is linearized overnight at 37oC with BsaI enzyme. The transcription reaction buffer (40 mM Tris-HCl, pH 7.7, 0.01% Triton X-100, 5 mM DTT) was supplemented with 5 mM from each NTP, 24 mM MgCl2, 1 U/ml Inorganic Pyrophosphatase from baker’s yeast, 200 μM FUS NTD (if added) and 0.3 μM T7 RNA polymerase. Finally, highly concentrated agarose (1.5 % wt/v) was added to achieve 0.5 % final concentration and transferred quickly, while liquid, to a 3 mm NMR tube with long glass pipette.

### Theoretical model of FUS maturation in coexisting phases

We devised a simple model of FUS maturation in coexisting phases that accounts for protein-solvent interactions. The model considers an incompressible, ternary mixture of monomers, aggregates, and solvent with volume fractions *ϕ*_1_, *ϕ*_2_ and *ϕ_s_* respectively. Monomers and aggregates can undergo phase separation as described by the Flory Huggins free energy density *f* [56]–[58]

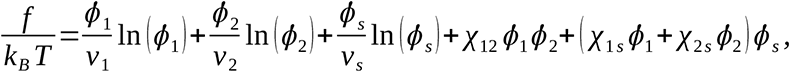

where *k _B_T* denotes thermal energy, *v_i_* is the molecular volume of component *i* (i=1,2,s), and *χ_ij_* are the Flory-Huggins interaction parameters describing the effective interaction between components *i* and *j* (monomer-solvent, aggregates-solvent, monomer-aggregates). Phase equilibrium between the droplet (I) and dilute (II) phases results from solving the phase-coexistence conditions:

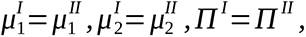

where *μ_i_* is the chemical potential of component *i*, *Π* is the osmotic pressure, and the superscript I/II indicates the dense/dilute phase, respectively.

In addition to undergoing phase separation, the monomers can aggregate. In each phase, we describe this process as a transition between monomeric and aggregated states via the reaction scheme *ϕ*_1_ *⇌ ϕ*_2_ . We note that the model can in principle be generalized to account for more complex aggregation events, including primary nucleation, elongation, fragmentation, or secondary nucleation [59], but further studies are required to understand the relative contribution of these aggregation steps to the overall aggregation process in droplets. We assume that the solvent is non-reactive and that the forward (*k*_1_) and backward (*k*_2_) rates are phase independent (i.e. they are identical in the dense and dilute phases). Under these conditions, we obtain the following kinetic equation for the average volume fractions of monomers and aggregates, 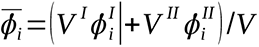 [17]:

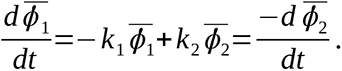

We consider the situation when phase separation is much faster than aggregation. Under these conditions, phase equilibrium is established almost instantly during aggregation. Therefore, we can solve the aggregation kinetics for the average monomer and aggregate volume fractions starting from a monomeric solution

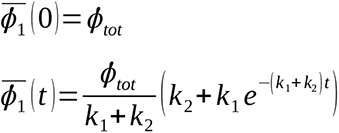

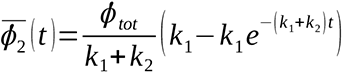

and at every time point during aggregation calculate the phase-separation equilibrium of the resulting monomer/aggregate mixture using phase coexistence conditions. This construction allows us to follow the time evolution of the monomer, aggregate and solvent concentrations in the dense (I) and dilute (II) phases, yielding the plots in Fig. S2.

To generate the plots, we used the following parameters: *v*_1_=*v*_2_=*v_s_*, *χ*_12_=0, *ϕ*_1_(0)=*ϕ_tot_* =0.55, *ϕ*_2_(0)=0, *χ*_1*s*_=3, *χ*_2*s*_=2.2, *k*_1_ =0.2*days*^−1^, *k*_2_ =0.1*days*^−1^ (Fig. S2b) and *v*_1_=*v*_2_=*v_s_*, *χ*_12_=0, *ϕ*_1_(0)=*ϕ_tot_* =0.55, *ϕ*_2_(0)=0, *χ*_1 *s*_=2.2, *χ*_2 *s*_=3, *k*_1_ =0.1*days*^−1^, *k*_2_ =0.05 *days*^−1^ (Fig. S2c).

### Microaspiration

The protein stock concentration for this experiment was 2 mM. The droplets were prepared by dilution to a final concentration of 100 μM with buffer without agarose (30 mM Hepes, 200 mM KCl, pH 7.3) and matured at room temperature over a period of a month. The micropipette with a 5 μm tip for this experiment was treated with BSA both on the inside and outside to prevent any clogging or adhesion of the condensed phase to the glass. RNA polyU purchased from SigmaAldrich was solubilized in the above buffer at stock concentration of 2 mg/ml. PolyU was added at a final concentration of 0.1 mg/ml. An external pump controlled the applied pressure of the micropipette (ΔP = 820 Pa) and a bright field microscope was used to visualize the microaspiration.

### Electron microscopy

A FUS NTD biphasic sample was matured for six months at r.t. inside a 3 mm NMR tube sealed by nail polish. The agarose hydrogel was extracted from the NMR tube by breaking the bottom of the tube. A 2 mm piece of hydrogel was cut into multiple small pieces, and subsequently, 50 μL dilution buffer (30 mM HEPES, 200 mM KCl, pH 7.3) was added. The sample was sonicated for 30 minutes at room temperature. Continuous carbon-supported copper grids (Quantifoil, Cu 300) were glow discharged (PELCO easiGlow, Ted Pella, negative, 25 mA, 30 s). After that, 3 µl of the sample was applied to the grids and incubated for 1 minute at r.t. The grids were then blot dried, washed with two drops of distilled water, and stained with two drops of 1% uranyl acetate for 30 seconds. Micrographs were acquired using a Tecnai F20 (Thermo Fisher Scientific) microscope operated at 200 kV and equipped with Falcon II camera, at 62000x magnification (pixel size: 1.7Å/pix) at a total dose of about 50 e^-^. The targeted defocus for the data acquisition was set to -3 µm.

### Preparation of stabilized FUS droplets for Coherent anti-Stokes Raman Scattering (CARS) imaging

An agarose solution 0.35%(wt/vol) of ultra-low melt agarose (Sigma) in dilution buffer (30 mM HEPES, 200 mM KCl, pH 7.3) was made in a 15 mL conical tube. The tube was then placed in a Thermomixer (Eppendorf) operating at 90 °C, shaking at 300 rpm until the solution was clear. This stock solution was used within two days.

To make samples, we used Grace Bio-Labs SecureSeal® Imaging spacers as gaskets on standard glass slides. The solution of agarose was placed in the Thermomixer and heated to 90 °C at 300rpm for at least 10 minutes to ensure it was warm. The protective films of the gaskets were removed, and the gasket was stuck to a glass slide. 18µL of warmed agarose solution was placed into the free space in the center of gasket, and 2µL of FUS stock (1.2 mM in storage buffer) was immediately added to this solution and very gently pipetted. The formation of droplets was observed by initially observing a whitish ring that diffused radially outward after the addition of the FUS stock to the agarose solution. A coverslip was then placed on top of the gasket, pressed on the sides, and the sample was sealed with nail polish. These samples were stable for more than three months, judging by the lack of obvious evaporation in the sealed sample.

As-prepared samples were allowed to sit for two hours before measuring, which are termed "fresh” samples for the CARS studies in this work. Samples were measured, then stored in a drawer in the lab until the next measurement, e.g., 1 month later.

### Coherent anti-Stokes Raman Scattering (CARS) imaging of stabilized FUS droplets

For the CARS measurements, samples were removed from the cabinet drawer and placed on the microscope. Our broadband coherent anti-Stokes Raman scattering (BCARS) microscope has been described elsewhere, and its application for liquid droplets has also been previously presented [60], [61]. Briefly, the pump/probe and Stokes pulses are generated in a dual-output, sub-nanosecond laser source (CARS-SM-30, Leukos), spatially and temporally overlapped at the sample plane of an inverted microscope (Eclipse Ti-U, Nikon), and tightly focused on the sample with a 100X, 0.85 NA air objective (LCPlan N, Olympus). The BCARS signal is filtered from the excitation pulses and focused onto the slit of a spectrograph (Shamrock 303i, Andor), which disperses the spectral components on a cooled CCD camera (Newport DU920P-BR-DD, Andor). Samples were mounted with the cover slip facing the objective. The samples were then raster scanned by moving a piezo stage (Nano-PDQ 375 HS, Mad City Labs), and the data acquisition was controlled via interface software in LabView 2015 (National Instruments).

Collected hyperspectral data were processed afterward in IgorPro (Wavemetrics) to extract the Raman-like spectra. The Raman-like spectra were obtained by phase-retrieval *via* a modified Kramers-Kronig transform using the surrounding agarose solution as the non-resonant reference [62]. The remaining error phase was removed using a Savitzky-Golay filter with a 2nd-order polynomial and window size 400 cm ^-1^, producing the Raman-like spectra for this work.

### Solid-state NMR spectroscopy

FUS NTD uniformly ^13^C and ^15^N labelled protein was mixed in 1:10 ratio with dilution buffer without agarose (30 mM HEPES, 200 mM KCl, pH 7.3) to form droplets. The droplets were centrifuged (25 °C, 10000 *g*, 10 min), resulting in circa 30 μL sedimented droplets and the top dilute phase was removed. The droplet phase was resuspended in 50 μL agarose containing dilution buffer and transferred quickly to the NMR rotor.

Solid-state NMR spectra were recorded at 20.0 T static magnetic-field strength in a 3.2 mm Bruker “Efree” probe [63]. The MAS frequency for all the experiments was set to 17 kHz. All spectra were processed with the software topspin (version 4.1.3, Bruker Biospin). The 2D spectra were processed with a shifted (DARR monophasic: 2.5, NCA/NCO monophasic: 3, DARR biphasic: 2) squared cosine apodization function and automated baseline correction in the indirect and direct dimensions. The sample temperature was set to 278 K [64]. All spectra were analyzed with the software CcpNmr (version 2.4.2) and referenced to 2,2-dimethyl-2-silapentane-5-sulfonate (DSS) [65], [66]. The experimental parameters used are summarized in Table S2.

### Analysis of real-time solid-state NMR kinetics

The kinetic analysis of the time-dependent intensities from the 1D spectra of biphasic and monophasic FUS was carried out by manually extracting individual signal-to-noise values of some representative peaks (Fig 2a,b), as well as absolute integral values (Fig S10), of each spectrum in the time-dependent series via the build-in top-spin module SiNo (signal-to-noise calculator, intensity of a peak divided by the square of the noise intensity). The intensities of interest were then loaded, visualized and processed in MATLAB (version R2021b, Natick, Massachusetts: The MathWorks Inc.).

### Secondary structure chemical-shift predictions

Average secondary structure-dependent chemical shift values for threonines and serines and their associated standard deviations were taken from reference [36] and visualized on the 1D CP spectrum using a home-written MATLAB script (version R2021b, Natick, Massachusetts: The MathWorks Inc.). While 2D probability density distribution plots for the secondary structure chemical-shift statistics were estimated and visualized using the *pluqin* python package, for which the raw data were extracted from the PASCY/BMRB database [37], [38].

## Acknowledgements

This work was supported by the Swiss National Science Foundation with Sinergia grant no. CR-SII5_170976 and NCCR RNA & Disease. LE acknowledges EMBO for the long-term postdoctoral fellowship (LTF-388-2018). TW acknowledges support from the Deutsche Forschungsgemeinschaft (DFG, German Research Foundation, project number 455240421 and Heisenberg fellowship, project number 455238107) and an ETH research grant (ETH-43 17-2 for funding of JZ). TW thanks the Max Planck Society for funding, as well as Prof. Dr. Beat H. Meier (ETH Zürich, Switzerland) for kindly providing measurement time for this project. This project was co-funded by SFB 1551 Project No. 464588647 of the Deutsche Forschungsgemeinschaft (DFG) to M.B. S.H.P. and Y.K. acknowledge support from the DFG project SPP 2191 #PA2526/3-1. We thank Dr. Marco E. Weber (ETH Zürich, Switzerland) for support with some solid-state NMR measurements and Dr. Y. Nikolaev for providing materials and protocol for the in vitro transcription reactions. In addition, we thank Prof. Eric Dufresne (ETH Zürich, Switzerland) for providing the microaspiration instrumentation and expertise.

## Author contributions

L.E. designed the project. L.E. performed and analyzed the liquid-state NMR experiments. E.B., J.Z. and T.W. performed and analyzed the solid-state NMR experiments. Y.K. and M.I. performed and analyzed the Raman and microaspiration experiments, respectively. M.E.P. performed the electron microscopy experiments with support of P.A. and D.B. T.C.T.M. developed the theoretical model of the solvent behavior. F.H.-T.A., S.H.P., M.B. and T.W. provided infrastructure, financial support and overall supervision of the project. L.E. and E.B. wrote the manuscript with support from all authors.

## Ethics declarations

The authors declare no competing interests.

## Data availability

Data supporting the findings of this study are available within the paper and its Supplementary Information.

## References

[1] S. Boeynaems et al., “Protein Phase Separation: A New Phase in Cell Biology,” Trends Cell Biol., vol. 28, no. 6, pp. 420–435, Jun. 2018, doi: 10.1016/J.TCB.2018.02.004.

[2] B. Wang et al., “Liquid-liquid phase separation in human health and diseases,” Signal Transduct. Target. Ther., vol. 6, no. 1, Dec. 2021, doi: 10.1038/S41392-021-00678-1.

[3] E. Gomes and J. Shorter, “The molecular language of membraneless organelles,” J. Biol. Chem., vol. 294, no. 18, pp. 7115–7127, May 2019, doi: 10.1074/JBC.TM118.001192.

[4] W. M. Babinchak and W. K. Surewicz, “Liquid-Liquid Phase Separation and Its Mechanistic Role in Pathological Protein Aggregation,” J. Mol. Biol., vol. 432, no. 7, pp. 1910–1925, Mar. 2020, doi: 10.1016/J.JMB.2020.03.004.

[5] J. Wang et al., “A Molecular Grammar Governing the Driving Forces for Phase Separation of Prion-like RNA Binding Proteins,” Cell, vol. 174, no. 3, pp. 688–699.e16, Jul. 2018, doi: 10.1016/J.CELL.2018.06.006/ATTACHMENT/E79432E0-30CF-4B59-8301-B8D9810EC81A/MMC6.XLSX.

[6] Y. Shin and C. P. Brangwynne, “Liquid phase condensation in cell physiology and disease,” Science, vol. 357, no. 6357. American Association for the Advancement of Science, Sep. 22, 2017, doi: 10.1126/science.aaf4382.

[7] J. Van Lindt et al., “A generic approach to study the kinetics of liquid–liquid phase separation under near-native conditions,” Commun. Biol. 2021 41, vol. 4, no. 1, pp. 1–8, Jan. 2021, doi: 10.1038/s42003-020-01596-8.

[8] E. Bentmann, M. Neumann, S. Tahirovic, R. Rodde, D. Dormann, and C. Haass, “Requirements for stress granule recruitment of fused in sarcoma (FUS) and TAR DNA-binding protein of 43 kDa (TDP-43),” J. Biol. Chem., vol. 287, no. 27, pp. 23079–23094, Jun. 2012, doi: 10.1074/JBC.M111.328757.

[9] A. Patel et al., “A Liquid-to-Solid Phase Transition of the ALS Protein FUS Accelerated by Disease Mutation,” Cell, vol. 162, no. 5, pp. 1066–1077, Aug. 2015, doi: 10.1016/J.CELL.2015.07.047.

[10] K. A. Burke, A. M. Janke, C. L. Rhine, and N. L. Fawzi, “Residue-by-Residue View of In Vitro FUS Granules that Bind the C-Terminal Domain of RNA Polymerase II,” Mol. Cell, vol. 60, no. 2, pp. 231–241, Oct. 2015, doi: 10.1016/j.molcel.2015.09.006.

[11] T. Murakami et al., “ALS/FTD Mutation-Induced Phase Transition of FUS Liquid Droplets and Reversible Hydrogels into Irreversible Hydrogels Impairs RNP Granule Function,” Neuron, vol. 88, no. 4, pp. 678– 690, Nov. 2015, doi: 10.1016/J.NEURON.2015.10.030.

[12] D. T. Murray et al., “Structure of FUS Protein Fibrils and Its Relevance to Self-Assembly and Phase Separation of Low-Complexity Domains,” Cell, vol. 171, no. 3, pp. 615–627.e16, Oct. 2017, doi: 10.1016/J.-CELL.2017.08.048.

[13] Y. Sun et al., “Molecular structure of an amyloid fibril formed by FUS low-complexity domain,” iScience, vol. 25, no. 1, p. 103701, Jan. 2022, doi: 10.1016/J.ISCI.2021.103701.

[14] R. F. Berkeley, M. Kashefi, and G. T. Debelouchina, “Real-time observation of structure and dynamics during the liquid-to-solid transition of FUS LC,” Biophys. J., vol. 120, no. 7, pp. 1276–1287, Apr. 2021, doi: 10.1016/J.BPJ.2021.02.008.

[15] L. Emmanouilidis et al., “NMR and EPR reveal a compaction of the RNA-binding protein FUS upon droplet formation,” Nat. Chem. Biol., 2021, doi: 10.1038/s41589-021-00752-3.

[16] J. P. Brady et al., “Structural and hydrodynamic properties of an intrinsically disordered region of a germ cell-specific protein on phase separation,” Proc. Natl. Acad. Sci. U. S. A., vol. 114, no. 39, pp. E8194– E8203, Sep. 2017, doi: 10.1073/pnas.1706197114.

[17] J. Bauermann, S. Laha, P. M. McCall, F. Jülicher, and C. A. Weber, “Chemical Kinetics and Mass Action in Coexisting Phases,” J. Am. Chem. Soc., vol. 144, no. 42, pp. 19294–19304, Oct. 2022, doi: 10.1021/JACS.2C06265.

[18] G. Bartolucci, T. C. T. Michaels, and C. A. Weber, “The interplay between molecular assembly and phase separation,” bioRxiv, p. 2023.04.18.537072, Apr. 2023, doi: 10.1101/2023.04.18.537072.

[19] L. Jawerth et al., “Protein condensates as aging Maxwell fluids,” Science (80-.)., vol. 370, no. 6522, pp. 1317–1323, Dec. 2020, doi: 10.1126/SCIENCE.AAW4951/SUPPL_FILE/AAW4951S3.MOV.

[20] J. C. Schwartz, X. Wang, E. R. Podell, and T. R. Cech, “RNA seeds higher-order assembly of FUS protein,” Cell Rep., vol. 5, no. 4, pp. 918–925, Nov. 2013, doi: 10.1016/J.CELREP.2013.11.017.

[21] I. Kwon et al., “Phosphorylation-regulated binding of RNA polymerase II to fibrous polymers of low-complexity domains,” Cell, vol. 155, no. 5, p. 1049, Nov. 2013, doi: 10.1016/J.CELL.2013.10.033.

[22] J. C. Schwartz, E. R. Podell, S. S. W. Han, J. D. Berry, K. C. Eggan, and T. R. Cech, “FUS is sequestered in nuclear aggregates in ALS patient fibroblasts,” Mol. Biol. Cell, vol. 25, no. 17, pp. 2571–2578, Sep. 2014, doi: 10.1091/MBC.E14-05-1007.

[23] Y. Nikolaev, N. Ripin, M. Soste, P. Picotti, D. Iber, and F. H. T. Allain, “Systems NMR: single-sample quantification of RNA, proteins and metabolites for biomolecular network analysis,” Nat. Methods, vol. 16, no. 8, pp. 743–749, Aug. 2019, doi: 10.1038/s41592-019-0495-7.

[24] S. Maharana et al., “RNA buffers the phase separation behavior of prion-like RNA-binding proteins,” Science, vol. 360, no. 6391, p. 918, May 2018, doi: 10.1126/SCIENCE.AAR7366.

[25] I. Matlahov and P. C. A. van der Wel, “Hidden motions and motion-induced invisibility: Dynamics-based spectral editing in solid-state NMR,” Methods, vol. 148, pp. 123–135, Sep. 2018, doi: 10.1016/J.YMETH.2018.04.015.

[26] A. B. Siemer, “Advances in studying protein disorder with solid-state NMR,” Solid State Nucl. Magn. Reson., vol. 106, p. 101643, Apr. 2020, doi: 10.1016/J.SSNMR.2020.101643.

[27] I. Bertini, G. Gallo, M. Korsak, C. Luchinat, J. Mao, and E. Ravera, “Formation Kinetics and Structural Features of Beta-Amyloid Aggregates by Sedimented Solute NMR,” ChemBioChem, vol. 14, no. 14, pp. 1891–1897, Sep. 2013, doi: 10.1002/CBIC.201300141.

[28] D. T. Murray and R. Tycko, “Sidechain hydrogen bonding interactions within amyloid-like fibrils formed by the low-complexity domain of FUS: Evidence from solid state nuclear magnetic resonance spectroscopy Graphical Abstract HHS Public Access,” Biochemistry, vol. 59, no. 4, pp. 364–378, 2020, doi: 10.1021/acs.biochem.9b00892.

[29] S. Elbaum-Garfinkle et al., “The disordered P granule protein LAF-1 drives phase separation into droplets with tunable viscosity and dynamics,” Proc. Natl. Acad. Sci. U. S. A., vol. 112, no. 23, pp. 7189–7194, Jun. 2015, doi: 10.1073/PNAS.1504822112/SUPPL_FILE/PNAS.201504822SI.PDF.

[30] S. Maharana et al., “RNA buffers the phase separation behavior of prion-like RNA binding proteins,” Science (80-.)., vol. 360, no. 6391, pp. 918–921, May 2018, doi: 10.1126/SCIENCE.AAR7366/SUPPL_FILE/AAR7366S4.MOV.

[31] K. Takegoshi, S. Nakamura, and T. Terao, “13C–13C polarization transfer by resonant interference recoupling under magic-angle spinning in solid-state NMR,” Chem. Phys. Lett., vol. 307, no. 5–6, pp. 295–302, Jul. 1999, doi: 10.1016/S0009-2614(99)00533-3.

[32] K. Takegoshi, S. Nakamura, and T. Terao, “13C–1H dipolar-assisted rotational resonance in magic-angle spinning NMR,” Chem. Phys. Lett., vol. 344, no. 5–6, pp. 631–637, Aug. 2001, doi: 10.1016/S0009-2614(01)00791-6.

[33] M. Baldus et al., “Cross polarization in the tilted frame: Assignment and spectral simplification in heteronuclear spin systems,” Mol. Phys., vol. 95, no. 6, pp. 1197–1207, 1998, doi: 10.1080/00268979809483251.

[34] S. Spera and A. Bax, “Empirical Correlation between Protein Backbone Conformation and Cα and Cβ 13C Nuclear Magnetic Resonance Chemical Shifts,” J. Am. Chem. Soc., vol. 113, no. 14, pp. 5490–5492, Jul. 1991, doi: 10.1021/JA00014A071/SUPPL_FILE/JA00014A071_SI_001.PDF.

[35] B. H. Meier and A. Böckmann, “The structure of fibrils from ‘misfolded’ proteins,” Curr. Opin. Struct. Biol., vol. 30, pp. 43–49, Feb. 2015, doi: 10.1016/J.SBI.2014.12.001.

[36] Wang and O. Jardetzky, “Probability-based protein secondary structure identification using combined NMR chemical-shift data,” Protein Sci., vol. 11, no. 4, pp. 852–861, Apr. 2002, doi: 10.1110/ps.3180102.

[37] K. J. Fritzsching, M. Hong, and K. Schmidt-Rohr, “Conformationally selective multidimensional chemical shift ranges in proteins from a PACSY database purged using intrinsic quality criteria,” J. Biomol. NMR, vol. 64, no. 2, pp. 115–130, Feb. 2016, doi: 10.1007/S10858-016-0013-5.

[38] K. J. Fritzsching, Y. Yang, K. Schmidt-Rohr, and M. Hong, “Practical use of chemical shift databases for protein solid-state NMR: 2D chemical shift maps and amino-acid assignment with secondary-structure information,” J. Biomol. NMR, vol. 56, no. 2, pp. 155–167, Jun. 2013, doi: 10.1007/S10858-013-9732-Z/TABLES/3.

[39] A. C. Murthy et al., “Molecular interactions contributing to FUS SYGQ LC/RGG phase separation and co-partitioning with RNA polymerase II heptads,” Nat. Struct. Mol. Biol., vol. 28, no. 11, p. 923, Nov. 2021, doi: 10.1038/S41594-021-00677-4.

[40] M. Callon et al., “Fast Magic-Angle-Spinning NMR Reveals the Evasive Hepatitis B Virus Capsid C-Terminal Domain,” Angew. Chem. Int. Ed. Engl., vol. 61, no. 32, Aug. 2022, doi: 10.1002/ANIE.202201083.

[41] F. Luo et al., “Atomic structures of FUS LC domain segments reveal bases for reversible amyloid fibril formation,” Nat. Struct. Mol. Biol. 2018 254, vol. 25, no. 4, pp. 341–346, Apr. 2018, doi: 10.1038/s41594-018-0050-8.

[42] A. Garaizar et al., “Aging can transform single-component protein condensates into multiphase architectures,” Proc. Natl. Acad. Sci. U. S. A., vol. 119, no. 26, Jun. 2022, doi: 10.1073/PNAS.2119800119/-/DC-SUPPLEMENTAL.

[43] M. Feric et al., “Coexisting Liquid Phases Underlie Nucleolar Subcompartments,” Cell, vol. 165, no. 7, pp. 1686–1697, Jun. 2016, doi: 10.1016/J.CELL.2016.04.047.

[44] S. Chatterjee et al., “Reversible Kinetic Trapping of FUS Biomolecular Condensates,” Adv. Sci., vol. 9, no. 4, p. 2104247, Feb. 2022, doi: 10.1002/ADVS.202104247.

[45] M. Farag, S. R. Cohen, W. M. Borcherds, A. Bremer, T. Mittag, and R. V. Pappu, “Condensates formed by prion-like low-complexity domains have small-world network structures and interfaces defined by expanded conformations,” Nat. Commun. 2022 131, vol. 13, no. 1, pp. 1–15, Dec. 2022, doi: 10.1038/s41467-022-35370-7.

[46] W. P. Lipiński et al., “Biomolecular condensates can both accelerate and suppress aggregation of α-synuclein,” Sci. Adv., vol. 8, no. 48, Dec. 2022, doi: 10.1126/SCIADV.ABQ6495.

[47] D. V. Laurents, C. Stuani, D. Pantoja-Uceda, E. Buratti, and M. Mompeán, “Aromatic and aliphatic residues of the disordered region of TDP-43 are on a fast track for self-assembly,” Biochem. Biophys. Res. Commun., vol. 578, pp. 110–114, Nov. 2021, doi: 10.1016/J.BBRC.2021.09.040.

[48] N. Nuraje, K. Su, N. I. Yang, and H. Matsui, “Liquid/Liquid Interfacial Polymerization To Grow Single Crystalline Nanoneedles of Various Conducting Polymers,” ACS Nano, vol. 2, no. 3, p. 502, Mar. 2008, doi: 10.1021/NN7001536.

[49] K. Piradashvili, E. M. Alexandrino, F. R. Wurm, and K. Landfester, “Reactions and polymerizations at the liquid-liquid interface,” Chem. Rev., vol. 116, no. 4, pp. 2141–2169, Feb. 2016, doi: 10.1021/ACS.CHEM-REV.5B00567/ASSET/IMAGES/LARGE/CR-2015-00567G_0005.JPEG.

[50] F. Puvion-Dutilleul, J. P. Bachellerie, and E. Puvion, “Nucleolar organization of HeLa cells as studied by in situ hybridization,” Chromosoma, vol. 100, no. 6, pp. 395–409, Jul. 1991, doi: 10.1007/BF00337518.

[51] R. W. Yao et al., “Nascent Pre-rRNA Sorting via Phase Separation Drives the Assembly of Dense Fibrillar Components in the Human Nucleolus,” Mol. Cell, vol. 76, no. 5, pp. 767–783.e11, Dec. 2019, doi: 10.1016/J.MOLCEL.2019.08.014.

[52] D. L. J. Lafontaine, J. A. Riback, R. Bascetin, and C. P. Brangwynne, “The nucleolus as a multiphase liquid condensate,” Nature Reviews Molecular Cell Biology, vol. 22, no. 3. Nature Research, pp. 165–182, Mar. 01, 2021, doi: 10.1038/s41580-020-0272-6.

[53] H. Yu et al., “HSP70 chaperones RNA-free TDP-43 into anisotropic intranuclear liquid spherical shells,” Science (80-.)., vol. 371, no. 6529, Feb. 2021, doi: 10.1126/SCIENCE.ABB4309.

[54] M. Caputi and A. M. Zahler, “Determination of the RNA Binding Specificity of the Heterogeneous Nuclear Ribonucleoprotein (hnRNP) H/H′/F/2H9 Family,” J. Biol. Chem., vol. 276, no. 47, pp. 43850–43859, Nov. 2001, doi: 10.1074/JBC.M102861200.

[55] E. Michel, O. Duss, and F. H. T. Allain, “An integrated cell-free assay to study translation regulation by small bacterial noncoding RNAs,” Methods Mol. Biol., vol. 1737, pp. 177–195, 2018, doi: 10.1007/978-1-4939-7634-8_11/FIGURES/8.

[56] C. P. Brangwynne, P. Tompa, and R. V. Pappu, “Polymer physics of intracellular phase transitions,” Nat. Phys. 2015 1111, vol. 11, no. 11, pp. 899–904, Nov. 2015, doi: 10.1038/nphys3532.

[57] A. W. Fritsch et al., “Local thermodynamics govern formation and dissolution of Caenorhabditis elegans P granule condensates,” Proc. Natl. Acad. Sci. U. S. A., vol. 118, no. 37, Sep. 2021, doi: 10.1073/PNAS.2102772118/-/DCSUPPLEMENTAL.

[58] J. A. Riback et al., “Composition-dependent thermodynamics of intracellular phase separation,” Nat. 2020 5817807, vol. 581, no. 7807, pp. 209–214, May 2020, doi: 10.1038/s41586-020-2256-2.

[59] T. C. T. Michaels et al., “Chemical Kinetics for Bridging Molecular Mechanisms and Macroscopic Measurements of Amyloid Fibril Formation,” https://doi.org/10.1146/annurev-physchem-050317-021322, vol. 69, pp. 273–298, Apr. 2018, doi: 10.1146/ANNUREV-PHYSCHEM-050317-021322.

[60] A. C. Murthy et al., “Molecular interactions underlying liquid−liquid phase separation of the FUS low-complexity domain,” Nat. Struct. Mol. Biol. 2019 267, vol. 26, no. 7, pp. 637–648, Jul. 2019, doi: 10.1038/s41594-019-0250-x.

[61] N. Billecke et al., “Chemical imaging of lipid droplets in muscle tissues using hyperspectral coherent Raman microscopy,” Histochem Cell Biol, vol. 141, pp. 263–273, 2014, doi: 10.1007/s00418-013-1161-2.

[62] Y. Liu, Y. J. Lee, and M. T. Cicerone, “Broadband CARS spectral phase retrieval using a time-domain Kramers–Kronig transform,” Opt. Lett. Vol. 34, Issue 9, pp. 1363-1365, vol. 34, no. 9, pp. 1363–1365, May 2009, doi: 10.1364/OL.34.001363.

[63] P. L. Gor’kov, R. Witter, E. Y. Chekmenev, F. Nozirov, R. Fu, and W. W. Brey, “Low-E probe for 19F–1H NMR of dilute biological solids,” J. Magn. Reson., vol. 189, no. 2, pp. 182–189, Dec. 2007, doi: 10.1016/J.JMR.2007.09.008.

[64] A. Böckmann et al., “Characterization of different water pools in solid-state NMR protein samples,” J. Biomol. NMR, vol. 45, no. 3, pp. 319–327, Sep. 2009, doi: 10.1007/S10858-009-9374-3/FIGURES/7.

[65] R. Fogh et al., “The CCPN project: an interim report on a data model for the NMR community,” Nat. Struct. Biol. 2002 96, vol. 9, no. 6, pp. 416–418, 2002, doi: 10.1038/nsb0602-416.

[66] W. F. Vranken et al., “The CCPN data model for NMR spectroscopy: Development of a software pipeline,” Proteins Struct. Funct. Bioinforma., vol. 59, no. 4, pp. 687–696, Jun. 2005, doi: 10.1002/PROT.20449.

