## Supplementary Information for "A solid beta-sheet structure is formed at the surface of FUS liquid droplets during aging"

**Table 1**: Overview of solid-state NMR investigations performed on different FUS constructs.

| FUS sequence | Solid-state NMR investigations | Reference |
|---|---|---|
| 1-214<br>monophasic | Structure determination of the fibril core (39-95) | Murray et al., 2017 |
| 1-163<br>monophasic | Maturation kinetics, structural studies | Berkeley et al., 2021 |
| 1-267<br>mono- and biphasic | Maturation kinetics, structural studies | This work |

**Table S2**: Overview about experimental parameters of the performed solid-state NMR experiments. For more details about the used adiabatic CP steps and the tangential shapes used see[1].

| **Sample** | **FUS NTD (monophasic)** | **FUS NTD (monophasic)** | **FUS NTD (monophasic)** | **FUS NTD (monophasic)** | **FUS NTD (monophasic)** | **FUS NTD (monophasic)** |
|---|---|---|---|---|---|---|
| **Experiment** | **1D $^{13}$C CP** | **1D $^{13}$C INEPT** | **2D DARR 20 ms** | **1D $^{15}$N,$^{1}$H CP-MAS** | **2D NCA** | **2D NCO** |
| $\nu_r$ / kHz | 17 | 17 | 17 | 17 | 17 | 17 |
| $B_0$ / T | 20 | 20 | 20 | 20 | 20 | 20 |
| **Transfer I** | **HC-CP** | **HC-INEPT** | **HC-CP** | **HN-CP** | **HN-CP** | **HN-CP** |
| $\nu_1(^1H)$ / kHz | 60 | - | 60 | 60 | 60 | 60 |
| $\nu_1(X)$ / kHz | 41.6 | - | 38 | 47 | - | - |
| $\nu_1(Y)$ / kHz | - | - | - | - | 43 | 43 |
| Shape | Tangent $^1$H | - | Tangent $^1$H | Tangent $^1$H | Tangent $^1$H | Tangent $^1$H |
| $^{13}$C carrier / ppm | 100 | 100 | 100 | - | - | - |
| $^{15}$N carrier / ppm | - | - | - | 120 | 120 | 120 |
| CP contact time / ms | 0.6 | - | 0.6 | 1.2 | 1.2 | 1.2 |
| **Transfer II** | - | - | **DARR** | - | **NC-CP** | **NC-CP** |
| $\nu_1(^1H)$ / kHz | - | - | 17 | - | - | - |
| $\nu_1(X)$ / kHz | - | - | - | - | 6 | 6 |
| $\nu_1(Y)$ / kHz | - | - | - | - | 11 | 11 |
| $^{13}$C carrier/ ppm | - | - | 100 | - | 100 | 100 |
| CP contact time / ms | - | - | 20 | - | 6.5 | 6.0 |
| $t_1$ increments | 3072 | 16384 | 1536 | 3072 | 3072 | 3072 |
| Sweep width ($t_1$) / ppm | 100 | 100 | 100 | 100 | 117 | 117 |
| Acquisition time ($t_1$) / ms | 15.4 | 81.9 | 7.7 | 15.4 | 15.4 | 15.4 |
| $t_2$ increments | - | - | 3’072 | - | 192 | 192 |
| Sweep width ($t_2$) / ppm | - | - | 100 | - | 16 | 16 |
| Acquisition time ($t_2$) / ms | - | - | 15.4 | - | 0.7 | 0.7 |
| $^1$H Spinal-64$^a$ or WALTZ-64$^b$ decoupling / kHz | 90$^a$ | 5$^b$ | 90$^a$ | 90$^a$ | 90$^a$ | 90$^a$ |
| Interscan delay / s | 2 | 2 | 2.7 | 1.5 | 2.7 | 2.7 |
| Number of scans | 1024 | 1024 | 36 | 1024 | 112 | 160 |
| Measurement time / h | 0.5 | 0.5 | 42 | 0.4 | 16.5 | 23.5 |

**Table S2 continued.**

| **Sample** | **FUS NTD (biphasic)** | **FUS NTD (biphasic)** | **FUS NTD (biphasic)** |
|---|---|---|---|
| **Experiment** | **1D $^{13}$C CP** | **1D $^{13}$C INEPT** | **2D DARR 20 ms** |
| $v_r$ / kHz | 17 | 17 | 17 |
| $B_0$ / T | 20 | 20 | 20 |
| **Transfer I** | **HC-CP** | **HC-INEPT** | **HC-CP** |
| $v_1(^1H)$ / kHz | 60 | - | 60 |
| $v_1$(X) / kHz | 40.8 | - | 40.8 |
| Shape | Tangent $^1$H | - | Tangent $^1$H |
| $^{13}$C carrier / ppm | 100 | 100 | 100 |
| CP contact time / ms | 0.6 | - | 0.6 |
| **Transfer II** | - | - | **DARR** |
| $v_1(^1H)$ / kHz | - | - | 17 |
| $^{13}$C carrier/ ppm | - | - | 100 |
| CP contact time / ms | - | - | 20 |
| *t*1 increments | 3072 | 16384 | 2560 |
| Sweep width (*t*1) / ppm | 100 | 100 | 100 |
| Acquisition time (*t*1) / ms | 15.4 | 81.9 | 12.8 |
| *t*2 increments | - | - | 3’072 |
| Sweep width (*t*2) / ppm | - | - | 100 |
| Acquisition time (*t*2) / ms | - | - | 15.4 |
| $^1$H Spinal-64[a] or WALTZ-64[b] decoupling / kHz | 90[a] | 5[b] | 90[a] |
| Interscan delay / s | 2 | 2 | 2.5 |
| Number of scans | 2048 | 2048 | 24 |
| Measurement time / h | 1.1 | 1.1 | 43 |

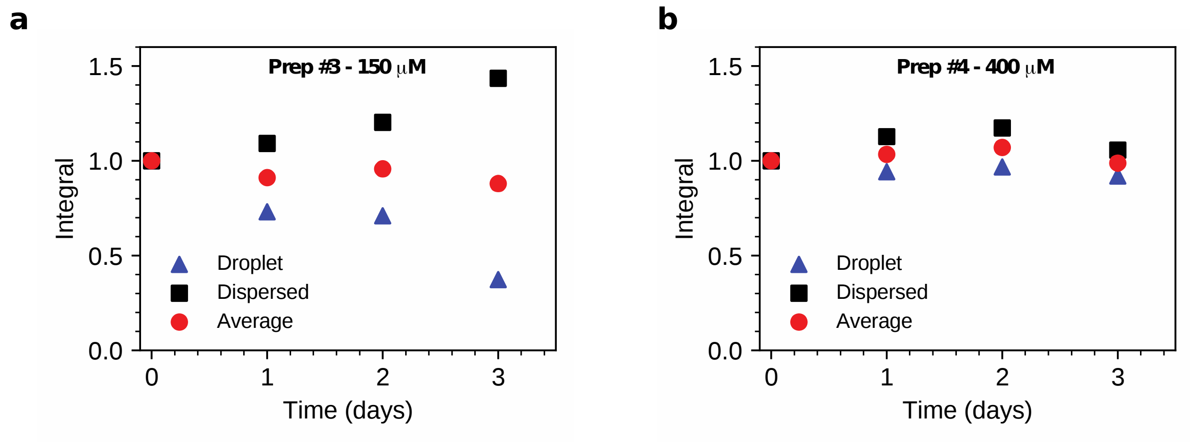


**Figure S1**: FUS fractions over time measured with different protein preparations and concentrations. a) At 150 µM of prep #3 droplet fraction simultaneously decays as the dispersed fraction increases. The two rates of change are equal as highlighted by their average which is stable at one. b) At 400 µM of prep #4 both droplet and dispersed fractions remained unaltered over time.

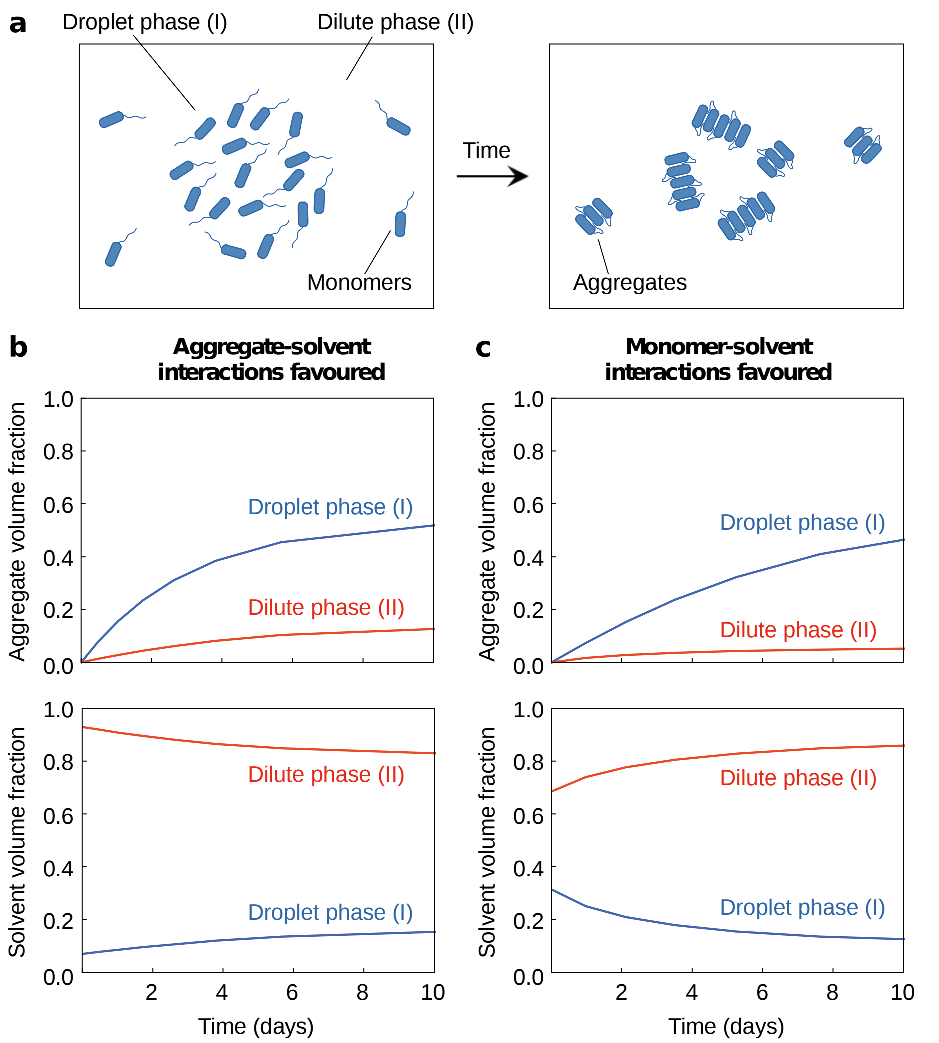


**Figure S2**: (a) Schematic representation of the model of FUS aggregation in coexisting phases. (b,c) Time evolution of aggregate and solvent volume fractions in the droplet (I) and dilute (II) phases for the case when (b) monomer-solvent interactions are disfavored over aggregate-solvent interactions ( $\chi_{1s} > \chi_{2s}$ ), or (c) aggregate-solvent interactions are disfavored over monomer-solvent interactions ( $\chi_{1s} < \chi_{2s}$ ). See Methods for parameters used in the plots.

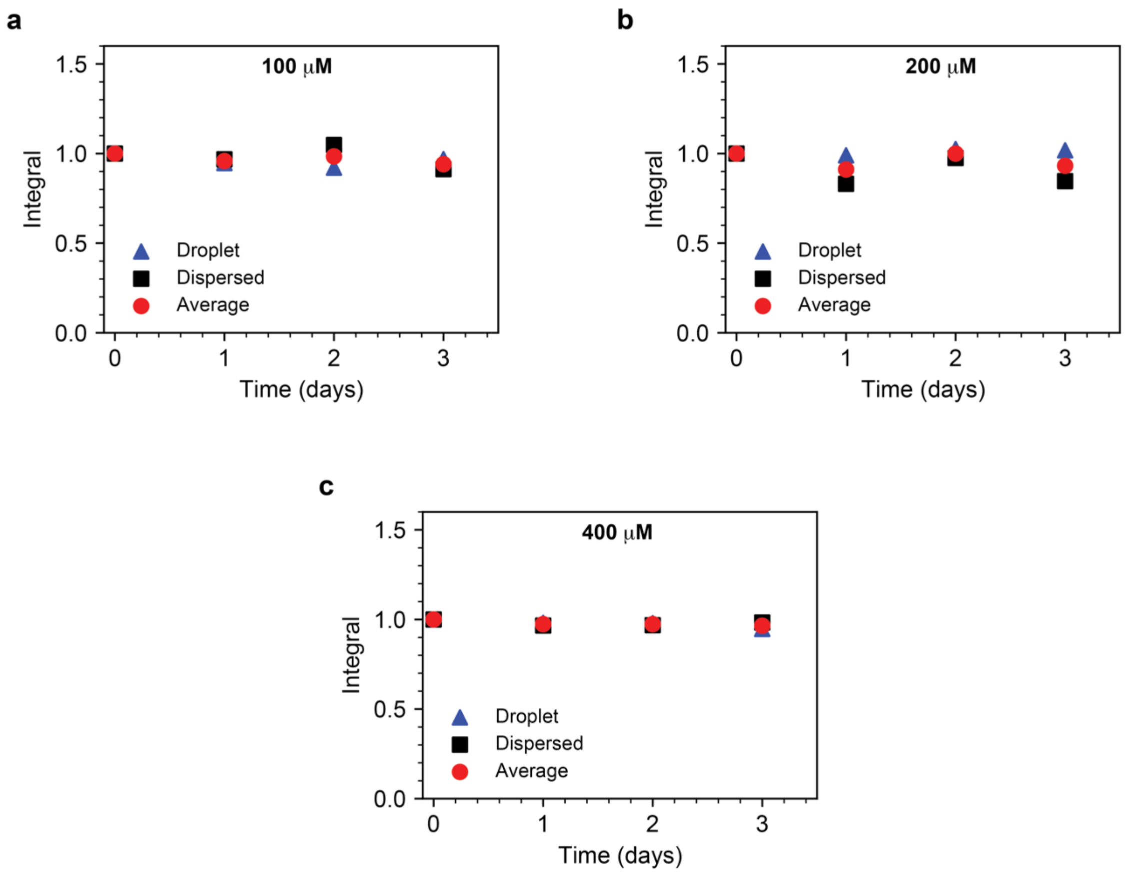


**Figure S3**: FUS fractions over time measured from different protein stocks at various concentrations. a, b, c) Both droplet and dispersed fractions of FUS did not change over the course of three days for all concentration measured (a: 100 μM, b: 200 μM, c: 400 μM).

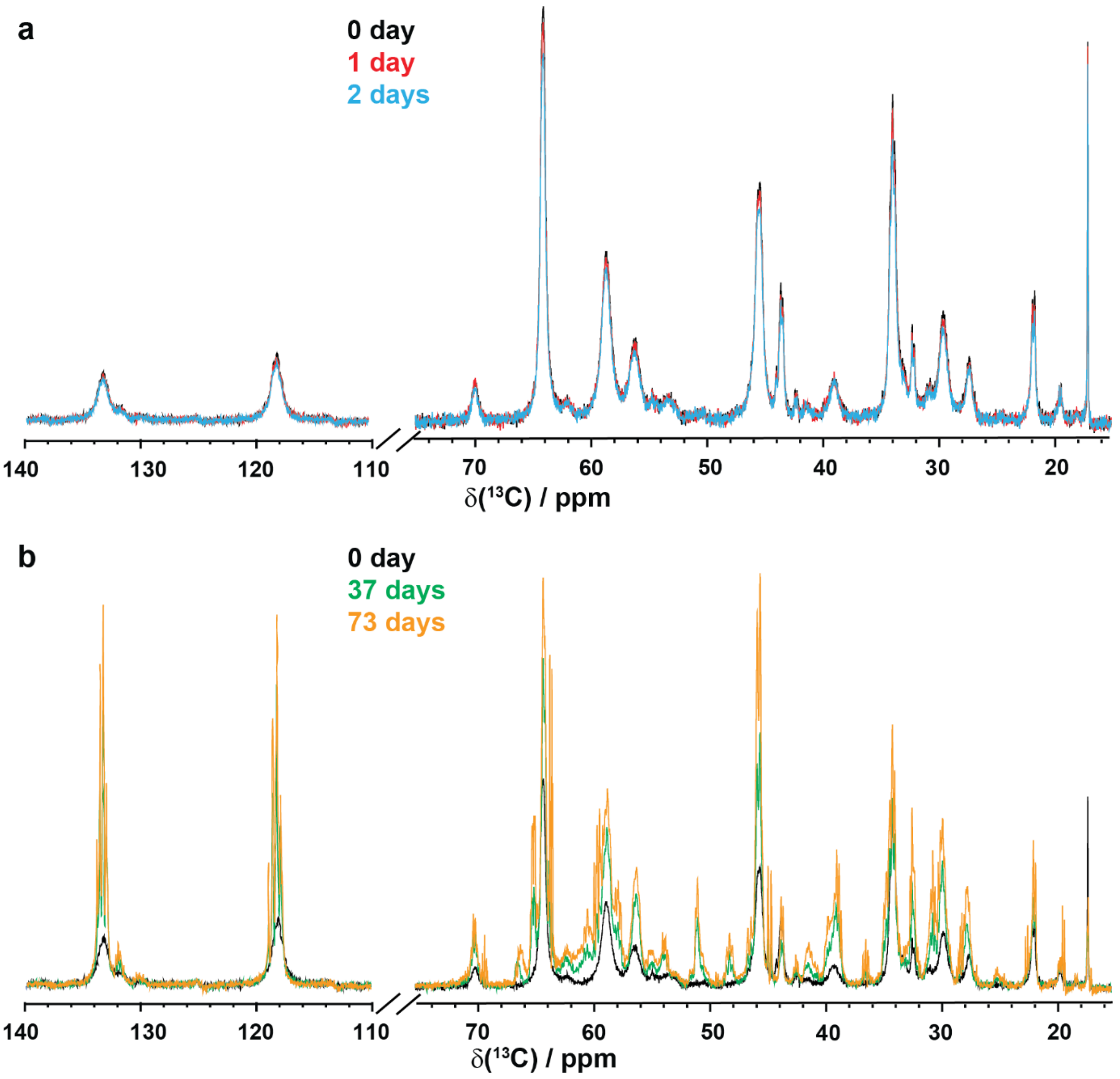


**Figure S4**: Comparison of 1D $^{13}C$-detected INEPT spectra for biphasic FUS-NTD liquid droplets recorded at various points in the maturation process. a) Almost no change in intensity is observed for the real-time INEPT signals up to two days of measurement after liquid droplet formation. b) Considerable increase in peak intensity, together with the appearance of *J*-resolved multiplet lines, can be appreciated already after 37 days. A further increase is observed after 73 days.

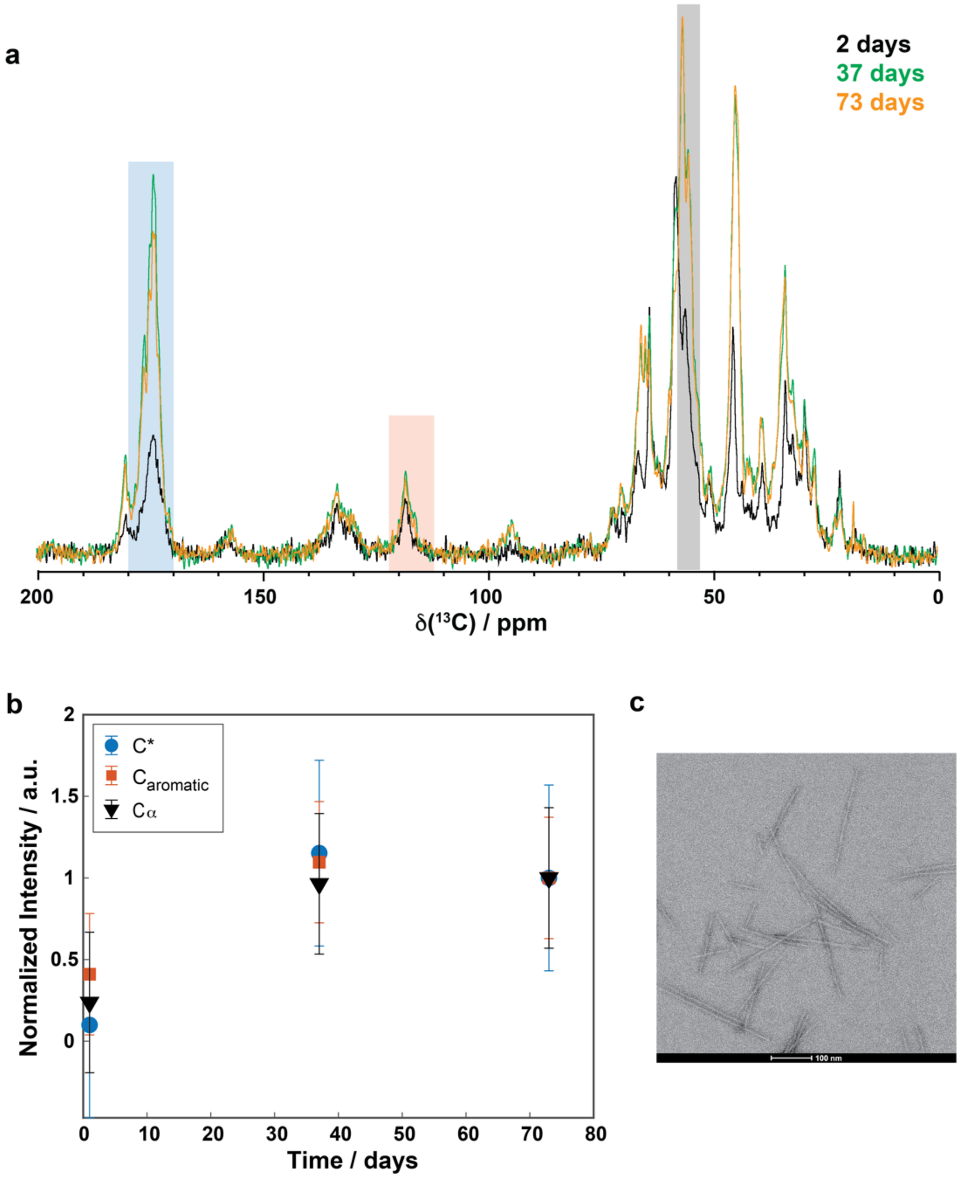


**Figure S5**: Maturation of biphasic FUS-NTD. a) Comparison of 1D $^{13}$C-detected CP spectra of biphasic FUS-NTD recorded at various points in the maturation process. The signal intensity stays rather constant over long-term maturation (i.e. from 37 to 73 days), indicating a plateau in the fibrilization process. b) The plateau in fibrilization is also confirmed by the kinetic analysis of the integrated intensities, which show small changes in intensity (within standard deviation) after 37 days and 73 days of storage. c) Electron-microscopy image of FUS fibrils taken on a biphasic sample matured for six months.

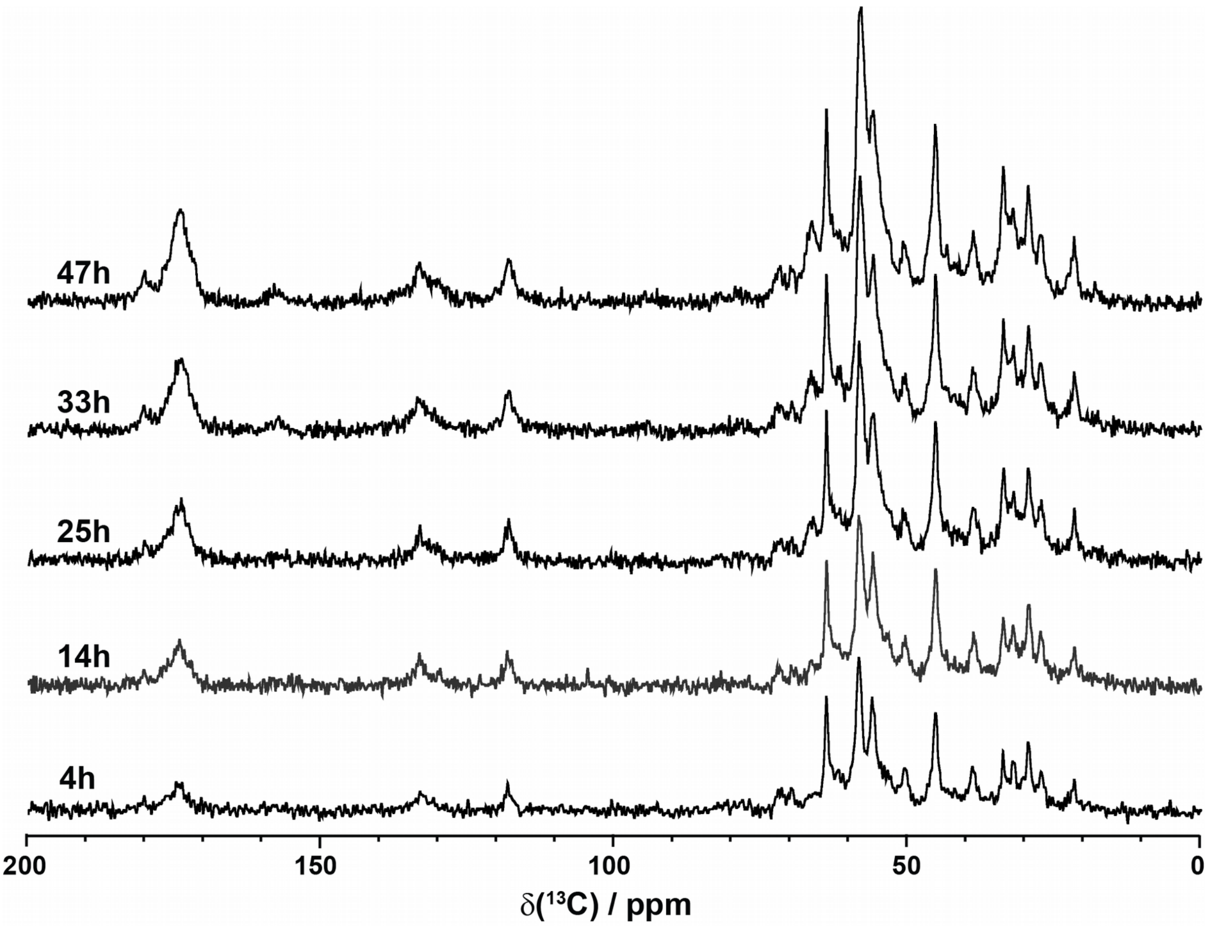


**Figure S6**: Overview of the time-dependent signal increase in the first two days of maturation for 1D $^{13}$C-detected CP spectra of biphasic FUS-NTD recorded at different time points. Consistent signal increase together with the absence of chemical-shift perturbations can be observed, supporting the conclusion of solid aggregate formation with a similar conformation.

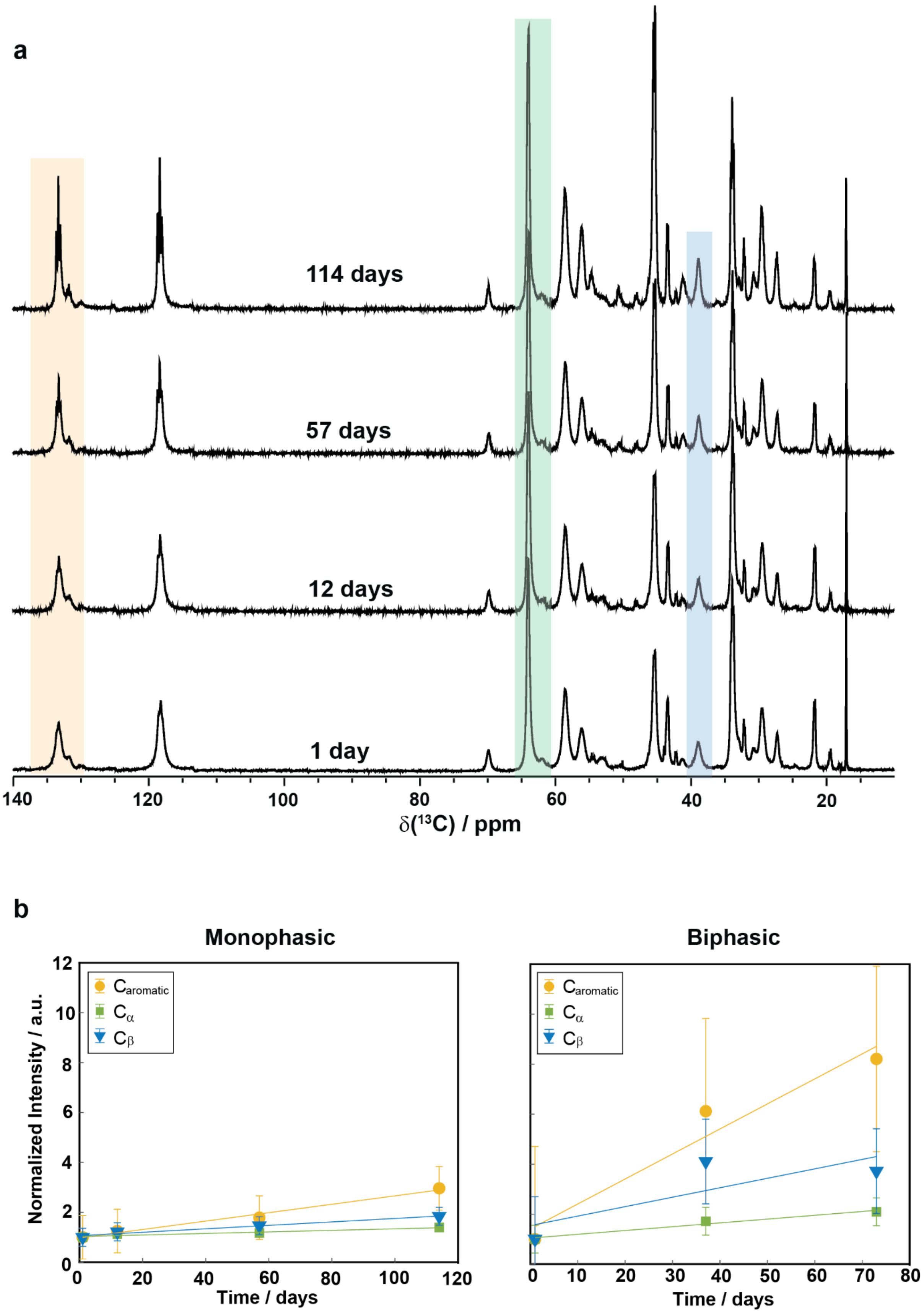


**Figure S7**: Overview of the signal increase in 1D $^{13}$C-detected INEPT spectra for monophasic FUS-NTD over 114 days of maturation. a) Stack plot of INEPT spectra recorded at four time points during the maturation period show an increase in signal intensity, accompanied by narrowing of the resonance lines). b) Comparison of the time dependence of the INEPT integrated intensities for the monophasic and biphasic sample indicates a smaller increase in intensity over time for the monophasic sample in comparison with the biphasic FUS-NTD.

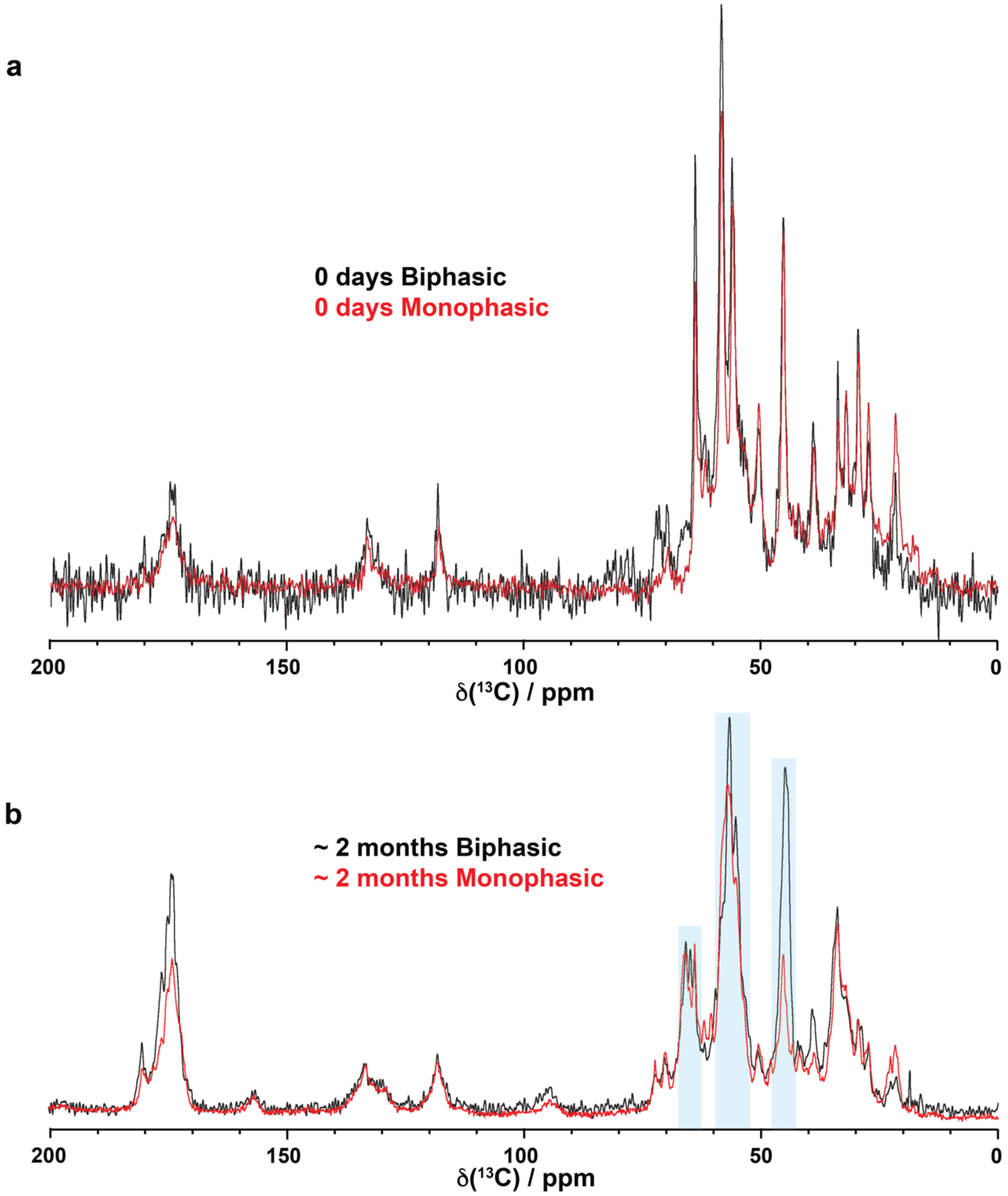


**Figure S8**: Comparison of 1D $^{13}C$ CP spectra of monophasic and biphasic FUS-NTD at the beginning of the maturation period (a) and after two months of storage at room temperature (b). a) The spectra look rather similar, albeit the threonine regions (~70 ppm) differ. b) After two months of maturation, notable differences can be observed, especially in the glycine region (~46 ppm) and carbonyl region for which a much more intense peak is observed for the biphasic sample.

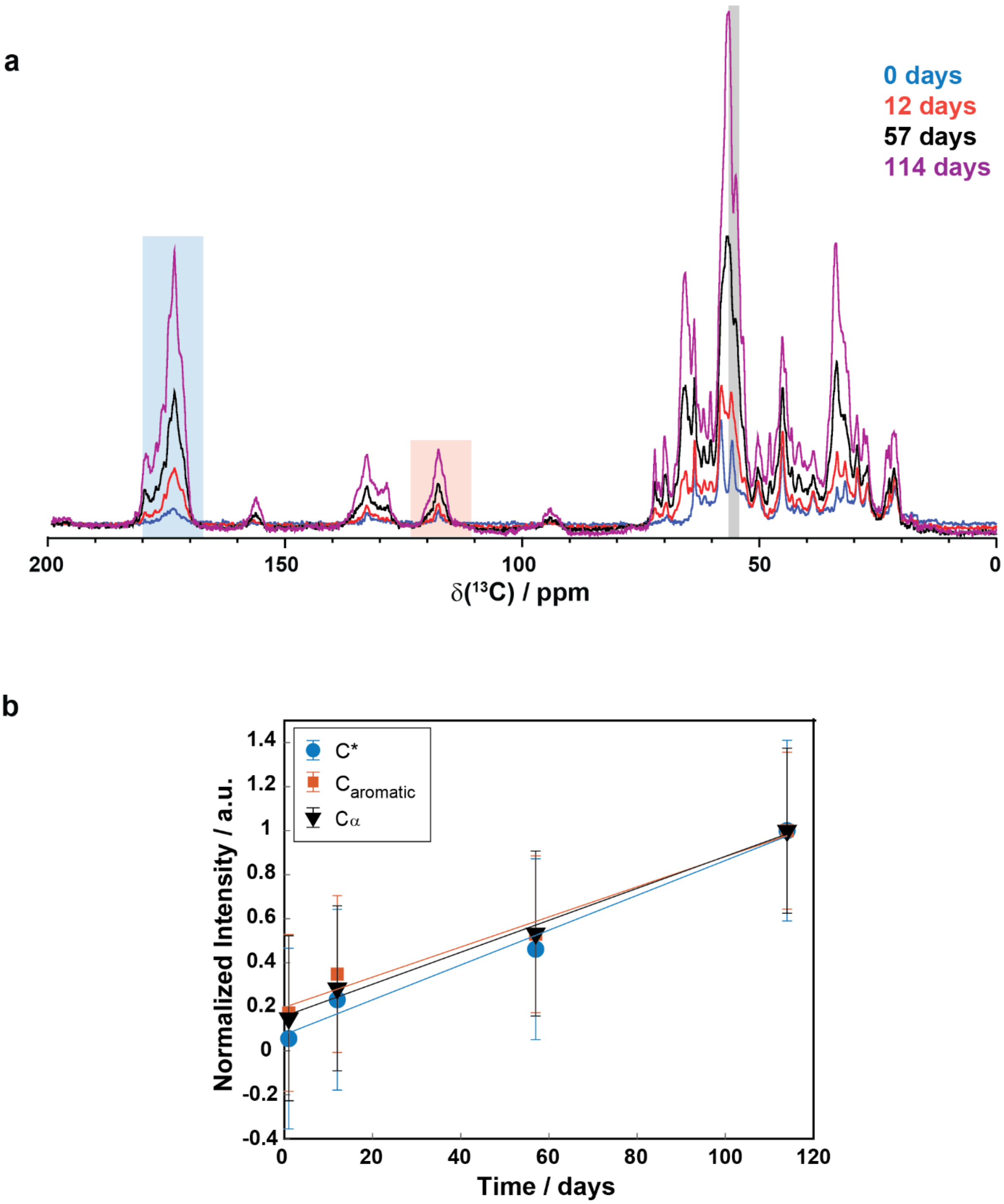


**Figure S9**: Maturation of monophasic FUS-NTD. a) Comparison of 1D $^{13}$C-detected CP spectra of monophasic FUS-NTD recorded at various points in the maturation process. Differently from the biphasic sample, the signal intensity increases consistently over the whole time up to 114 days. b) The kinetic analysis of the integrated intensities shows a rather linear increase over time. Spectral regions highlighted in a) were used for the analysis.

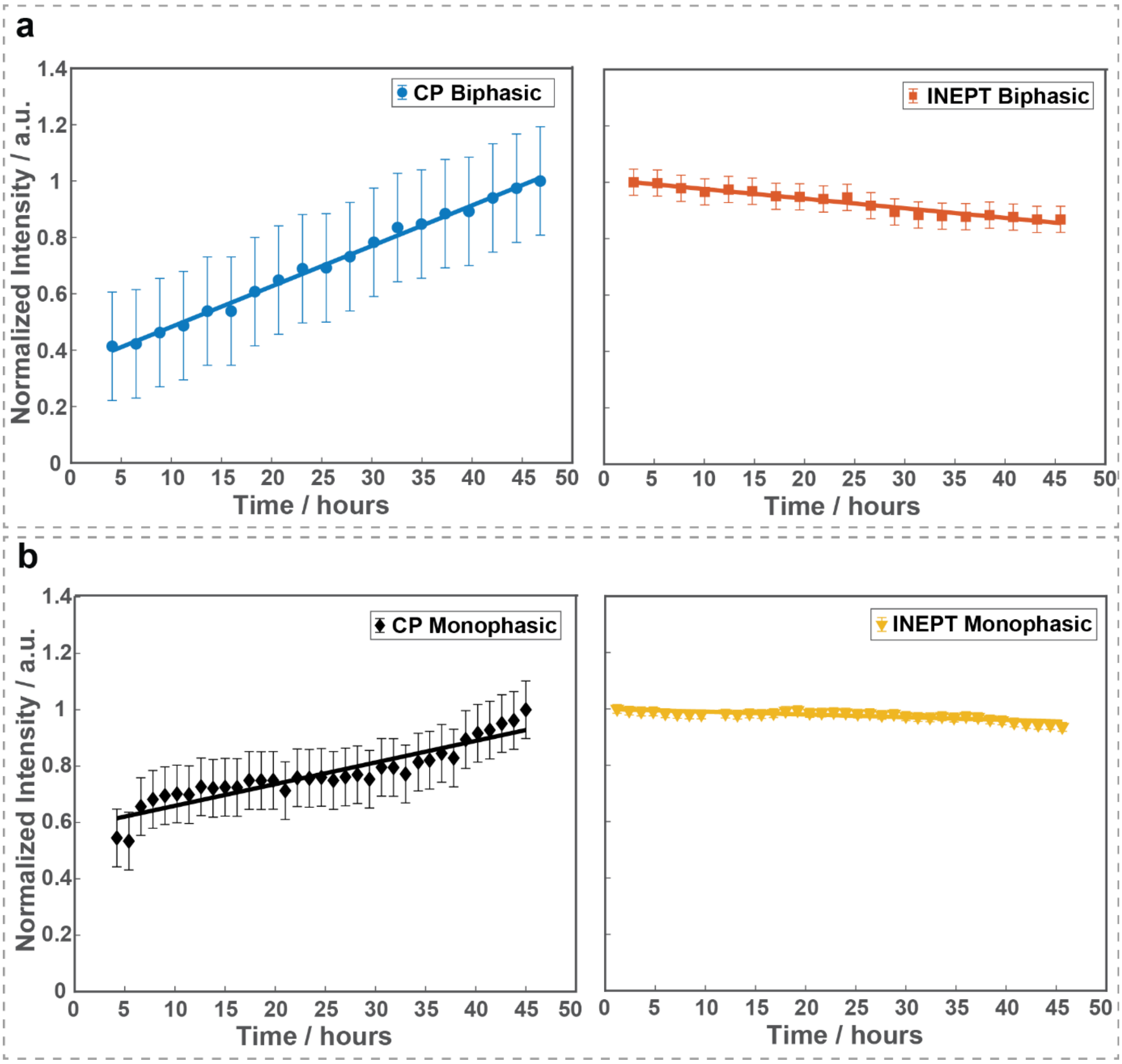


**Figure S10**: a) Time-dependent intensity changes for the absolute spectral integrals during two days of maturation for $^{13}C$,$^{1}H$ CP-MAS and $^{13}C$,$^{1}H$ INEPT spectra of biphasic FUS. The intensity of the total CP spectrum recorded after two days was normalized to one. A linear regression is shown (straight lines) with slopes of 0.346 a.u./days (CP), -0.082 a.u./days (INEPT). b) Time-dependent intensity changes for the absolute spectral integrals during two days of maturation for $^{13}C$,$^{1}H$ CP-MAS and $^{13}C$,$^{1}H$ INEPT spectra of monophasic FUS. The intensity of the total CP spectrum recorded after two days was normalized to one. A linear regression is shown (straight lines) with slopes of 0.214 a.u./days (CP), -0.022 a.u./days (INEPT).

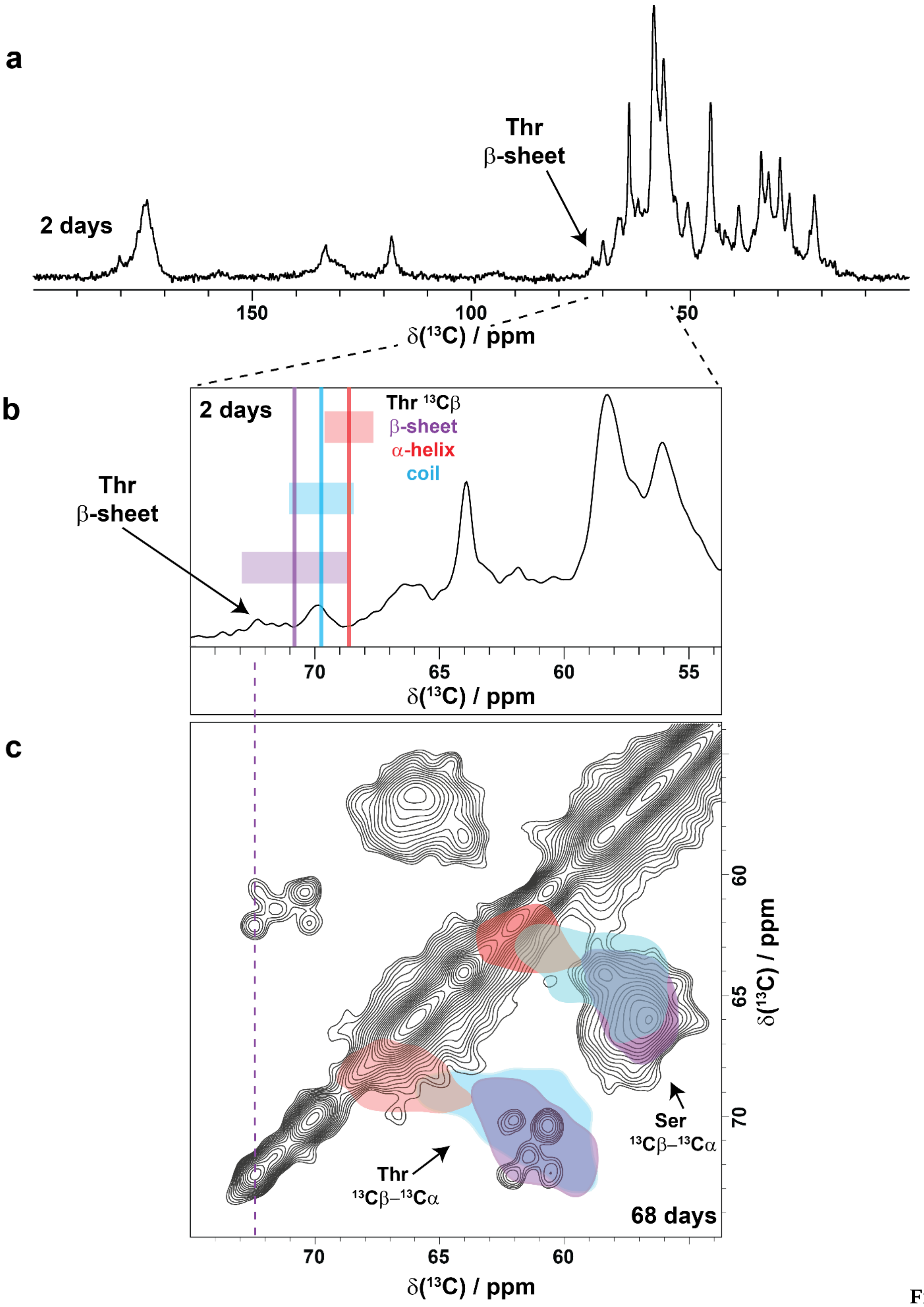


**Figure S11**: Secondary-structure chemical shift statistics indicates β-sheet formation. a) 1D $^{13}$C CP spectrum of monophasic FUS NTD recorded 2 days after preparation. b) Zoom into the $^{13}$C Cα/Cβ region of the spectrum with threonine Cβ chemical shift statistics plotted on top. c) Zoom into the 2D $^{13}$C-$^{13}$C DARR spectrum of monophasic maturated (68 days) FUS-NTD together with chemical-shift predictions for the well-resolved threonine and serine resonances. The details for the secondary-structure predictions of $^{13}$C Cα and Cβ chemical-shift values are reported in the material and methods section. The three types of secondary structure elements are highlighted with different colors.

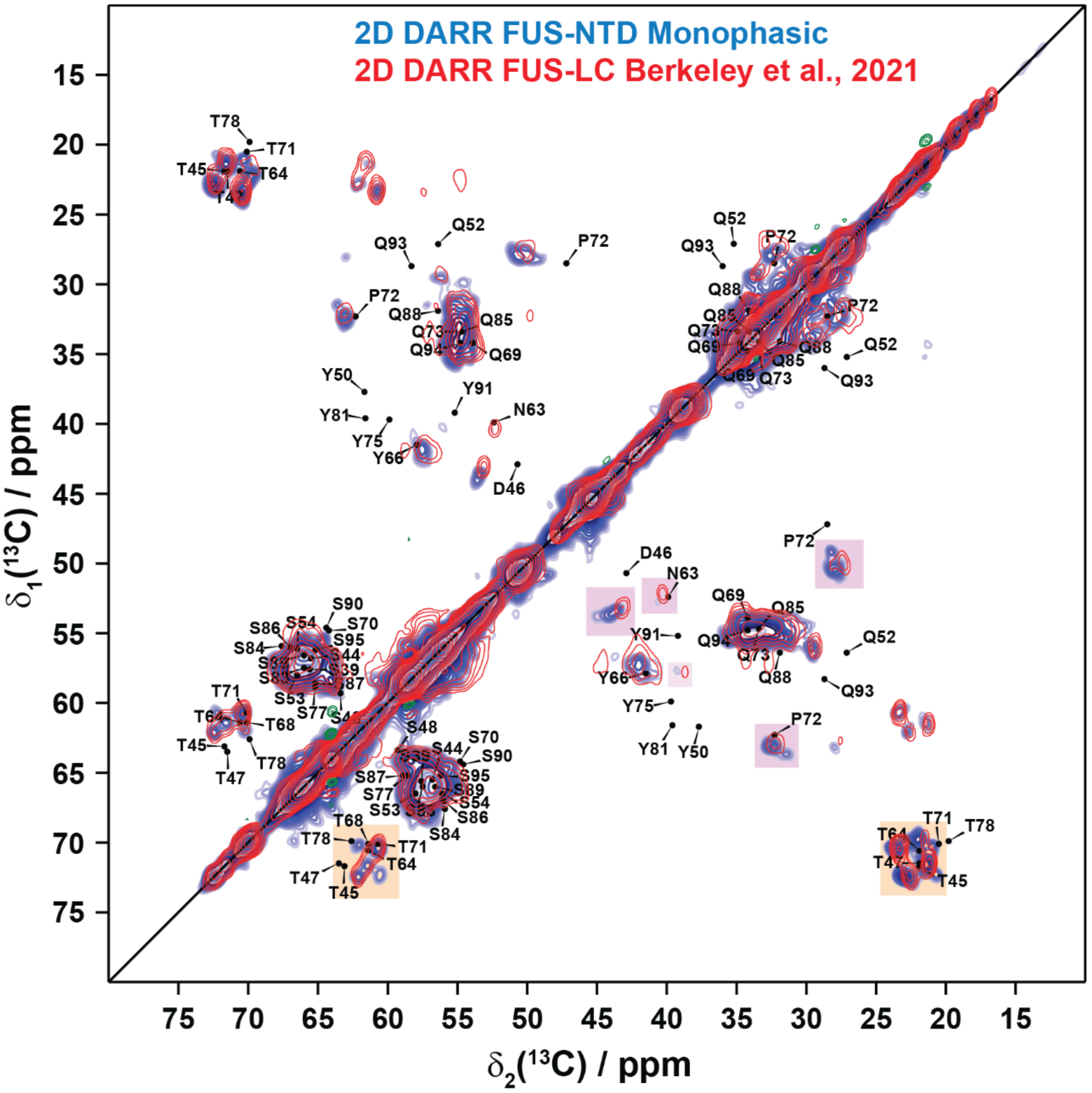


**Figure S12**: Comparison between the 2D DARR spectra for the maturated monophasic FUS-NTD in blue with the 2D DARR previously reported by [3] in red. Black labels report on the back-predicted shifts for the FUS fibril core from [4]. In the figure, orange boxes represent additional resonances in our DARR spectrum, while purple squares indicate chemical-shift perturbations.

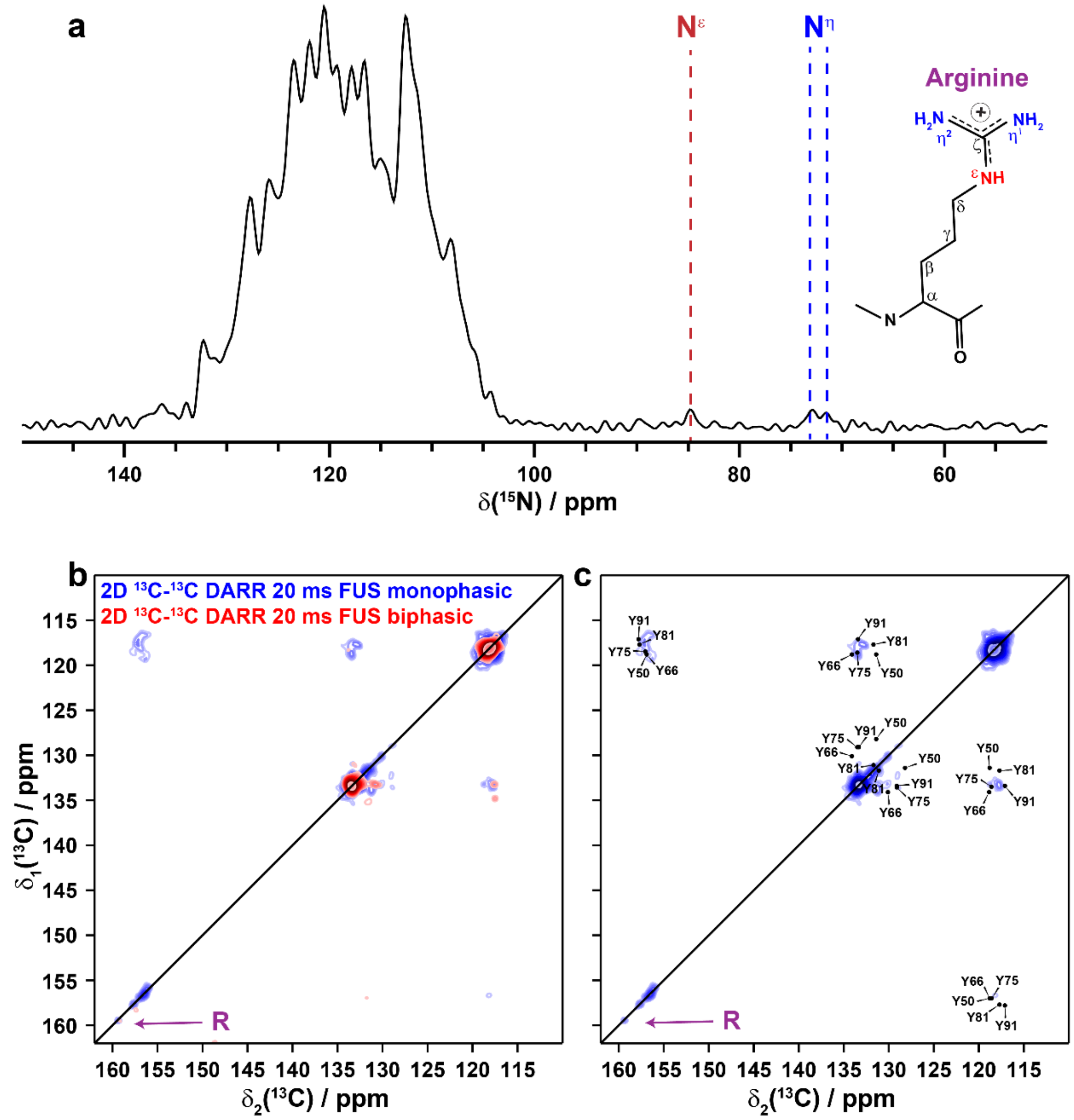


**Figure S13**: a) 1D $^{15}$N CP MAS spectrum of monophasic FUS NTD. The signals arising from the $N^{\varepsilon}$ and $N^{\eta}$ atoms of the arginine side chains are highlighted in red and blue respectively. b) Comparison of the aromatic regions of the 2D DARR spectra recorded on maturated mono- and biphasic FUS-NTD samples. In the monophasic sample, a Cζ arginine side chain diagonal peak highlighted by the purple arrow is observed. c) Aromatic region of the 2D DARR spectrum of the monophasic sample with back-predicted shifts for the FUS fibril core from [4].

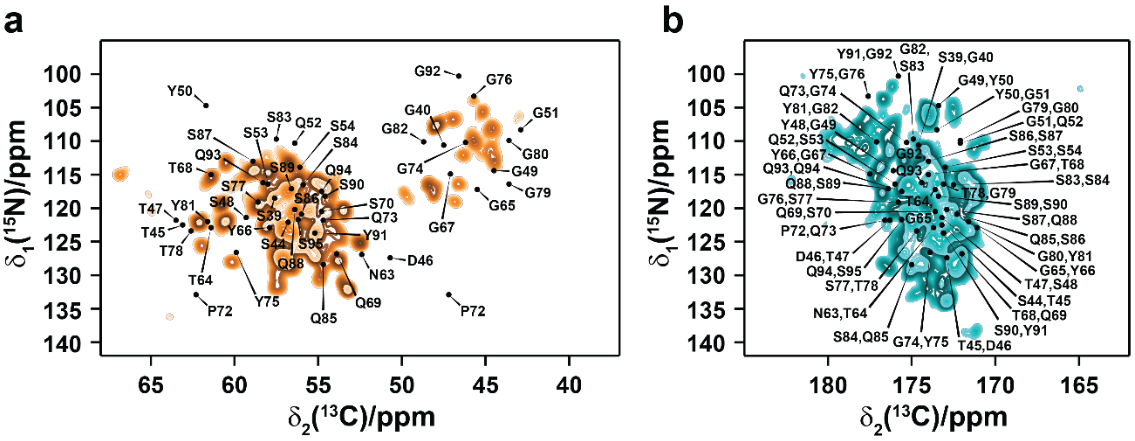

**Figure S14**: 2D NCA (a) and NCO (b) spectra of matured FUS-NTD liquid droplets (sample stored at 114 days at r.t.) shown also in Figure 2d of the main text. The assigned peaks are back-predicted from [4]. Both spectra were recorded at 20.0 T and 17 kHz MAS.

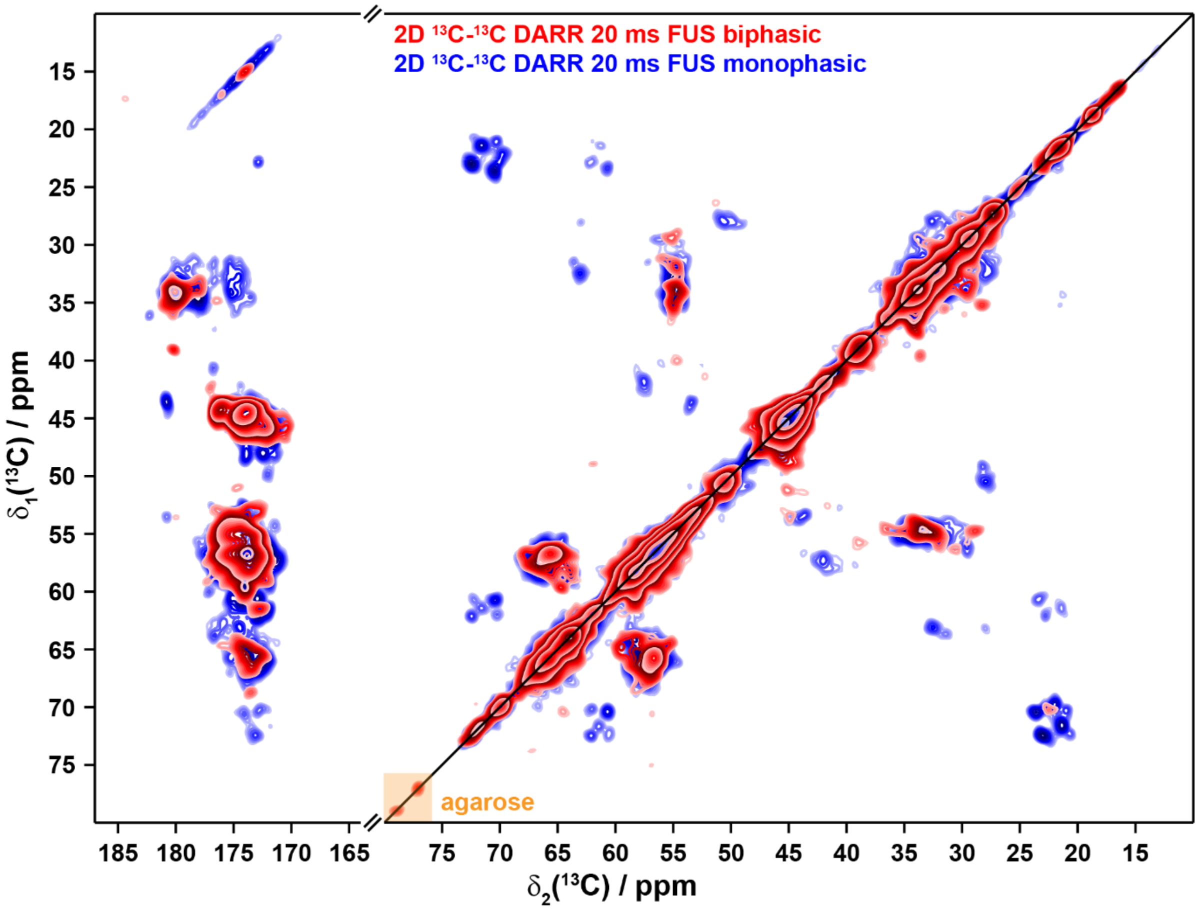


**Figure S15**: Comparison of 2D DARR spectra of the maturated mono- and biphasic FUS-NTD samples (68 days and 37 days of maturation). The orange box indicates $^{13}C$ signals originating from the agarose hydrogel matrix of the biphasic sample.

### Supplementary references


[1] a) S. Hediger, B. H. Meier, N. D. Kurur, G. Bodenhausen, R. R. Ernst, NMR cross polarization by adiabatic passage through the Hartmann—Hahn condition (APHH), *Chem. Phys. Lett.* **1994**, *223*, 283-288; b) S. Hediger, B. H. Meier, R. R. Ernst, Adiabatic passage Hartmann-Hahn cross polarization in NMR under magic angle sample spinning, *Chem. Phys. Lett.* **1995**, *240*, 449-456.

[2] Y. Wang, O. Jardetzky, Probability-based protein secondary structure identification using combined NMR chemical-shift data, *Protein Science : A Publication of the Protein Society* **2002**, *11*, 852-861.

[3] R. F. Berkeley, M. Kashefi, G. T. Debelouchina, Real-time observation of structure and dynamics during the liquid-to-solid transition of FUS LC, *Biophys. J.* **2021**, *120*, 1276-1287.

[4] D. T. Murray, M. Kato, Y. Lin, K. R. Thurber, I. Hung, S. L. McKnight, R. Tycko, Structure of FUS Protein Fibrils and Its Relevance to Self-Assembly and Phase Separation of Low-Complexity Domains, *Cell* **2017**, *171*, 615-627.e616.
